# Nuclear export of chimeric mRNAs depends on an lncRNA-triggered autoregulatory loop

**DOI:** 10.1101/426742

**Authors:** Zhen-Hua Chen, Zhan-Cheng Zeng, Tian-Qi Chen, Cai Han, Yu-Meng Sun, Wei Huang, Lin-Yu Sun, Ke Fang, Xue-Qun Luo, Wen-Tao Wang, Yue-Qin Chen

## Abstract

Aberrant chromosomal translocations leading to tumorigenesis have been ascribed to the heterogeneously oncogenic functions. However, how fusion transcripts exporting remains to be declared. Here, we showed that the nuclear speckle-specific long non coding RNA MALAT1 controls chimeric mRNA export processes and regulates myeloid progenitor cell differentiation in malignant hematopoiesis. We demonstrated that MALAT1 regulates chimeric mRNAs export in an m6A-dependent manner and thus controls hematopoietic cell differentiation. Specifically, reducing MALAT1 or m6A methyltransferases and the ‘reader’ YTHDC1 result in the universal retention of distinct oncogenic gene mRNAs in nucleus. Mechanically, MALAT1 hijacks both the chimeric mRNAs and fusion proteins in nuclear speckles during chromosomal translocations and mediates the colocalization of oncogenic fusion proteins with METTL14. MALAT1 and fusion protein complexes serve as a functional loading bridge for the interaction of chimeric mRNA and METTL14. This study demonstrated a universal mechanism of chimeric mRNA transport that involves lncRNA-fusion protein-m6A autoregulatory loop for controlling myeloid cell differentiation. Targeting the lncRNA-triggered autoregulatory loop to disrupt chimeric mRNA transport might represent a new common paradigm for treating blood malignancies.

## Introduction

Genomic alterations, particularly aberrant chromosomal translocations, are responsible for the onset of many types of cancers(Mertens et al, 2015; Mitelman et al, 2007), such as leukemia. For example, *PML-RARα* (t15; t17)(de The et al, 1991; Grignani et al, 1993), *MLL* fusions (t11)(Ayton & Cleary, 2001; Krivtsov & Armstrong, 2007; Meyer et al, 2009) and *AML1-ETO* (t8; t21)(Gardini et al, 2008; Li et al, 2016; Okumura et al, 2008) are typical oncogenic fusion genes that contribute to particular subtypes of leukemogenesis. How such rearrangements can lead to tumorigenesis has traditionally been explained by their ability to encode and express proteins; such proteins are commonly referred to as oncogenic ‘fusion proteins’(Mitelman et al, 2007; Racanicchi et al, 2005), indicating that the regulation of mRNA export of fusion proteins is important for oncogenic protein expression. However, how the resulting chimeric mRNA exports from the nucleus to the cytoplasm, as well as whether such alterations of the genome in the nucleus could also simultaneously impact the nuclear architecture, has been poorly characterized to date. Importantly, regulation of RNA exporting process has been shown to contribute to the drug-induced eradication of cancer cells(Harris, 2004). Thus, a greater understanding the mRNA export process of chromosomal translocations resulting in fusion genes is especially important for clinical therapeutic drug development(Biondi et al, 2000; Racanicchi et al, 2005).

It has been generally considered that most mRNA export is modulated by TAP-p15 heterodimers(Katahira, 2015; Katahira et al, 1999; Santos-Rosa et al, 1998), as well as a series of mRNA binding proteins and adaptor proteins(Hocine et al, 2010; Katahira, 2015). Recent studies have shown that the mRNA export process is also characterized by modifications of pre-mRNA, including N6-methyladenosine (m6A) modifications(Edupuganti et al, 2017; Geula et al, 2015; Zheng et al, 2013). The identified m6A methyltransferase complex comprised of methyltransferase-like 3 (METTL3), METTL14, and Wilms’ tumor 1-associating protein (WTAP) has been found to be specifically located in nuclear speckles(Liu et al, 2014), a nuclear structure related to mRNA export(Liu et al, 2014). In addition, nuclear noncoding RNAs (ncRNAs) are emerging as essential regulators of mRNA export^37^. The long noncoding RNA (lncRNA) NEAT1 is reportedly responsible for retaining double-stranded mRNAs in the nucleus (Chen & Carmichael, 2009; Fox & Lamond, 2010; Jiang et al, 2017; Scadden, 2009), suggesting that the mRNA export process is under complex regulatory control. Very interestingly, autoregulatory loops often provide precise control of the mRNA exporting process of specific genes that encode key proteins(Terenzio et al, 2018; Wu et al, 2015). For example, nuclear YRA1 autoregulates its own mRNA in trans, committing its pre-mRNA to the nuclear export process(Dong et al, 2007). Fusion protein partners, such as PML and AF4, are reported to have the capacity to bind to nascent RNA and their own mRNAs in the nucleus (Bitoun et al, 2007; Matera, 1999; Melko et al, 2011; Strudwick & Borden, 2002). The questions how fusion transcripts are exported from nucleus to cytoplasm and whether they are in potential autoregulation loop systems remains to be declared.

In this study, we performed genome-wide screening via RNA immunoprecipitation sequencing (RIP-seq) assay to assess fusion protein-associated RNAs in the nucleus. We discovered that MALAT1 directly interacted with several fusion proteins, including PML-RARA, MLL-AF9, MLL-ENL and AML1-ETO in nuclear speckles and affected the chimeric mRNA nucleocytoplasmic export and their protein production. We further showed that MALAT1 acted as a modulator that promotes fusion protein and m6A methyltransferase interactions, which in turn control the chimeric mRNA exporting process through the m6A reader YTHDC1. This study is the first to demonstrate that nuclear export of chimeric mRNAs depends on an lncRNA-triggered autoregulatory loop. The universal m6A-dependent autoregulatory mechanism of mRNA transport of distinct fusion proteins revealed in this study may provide a new therapeutic paradigm for treating a variety of leukemia subtypes.

## Results

### Both the mRNA and fusion proteins resulting from chromosomal rearranged genes were hijacked in nuclear speckles by MALAT1

To determine if there are any regulatory RNAs or other molecules that fusion proteins directly interact with, we used the PML-RARα fusion protein as a model, which is a specific t (15;17) chromosomal translocation resulting in the fusion of the promyelocytic gene (PML) and the retinoic acid receptor alpha gene (RARA) into the oncoprotein PML-RARA(de The et al, 1991; Grignani et al, 1993). We generated a stable NB4 cell line expressing FLAG-tagged PML-RARα using a lentivirus system in NB4 cells, and then performed RIP-seq assays to access the RNAs interacting with PML-RARα in cells. The specificity of the flag immunoprecipitation assay is shown in **Fig EV1A**. RIP-seq identified a number of regulatory noncoding transcripts and mRNA transcripts potentially interacting with PML-RARα (**Fig 1A**). Interestingly, MALAT1, a nuclear speckle-specific lncRNA, was identified as the most abundant ncRNA in deep sequencing data (**Fig 1A**). To further validate this direct interaction and to determine the interacting residues of MALAT1, we carried out MS2-trap assay(Yoon et al, 2012) as shown in **Fig 1B** using HA-tagged PML-RARα cDNA plasmid, which well expressed an endogenous fusion protein (**Fig EV1B**); we also generated three partial MALAT1 constructs (**Fig 1C**). The results confirmed the interaction between MALAT1 and PML-RARα; specifically, the MALAT1 3’ end section interacted with PML-RARα tightly much more than did the other sections (**Fig 1C**), consistent with previous findings that MALAT1 functions primarily through its triple helix structure(Brown et al, 2014; Brown et al, 2016; Liu et al, 2015).

**Figure 1.**
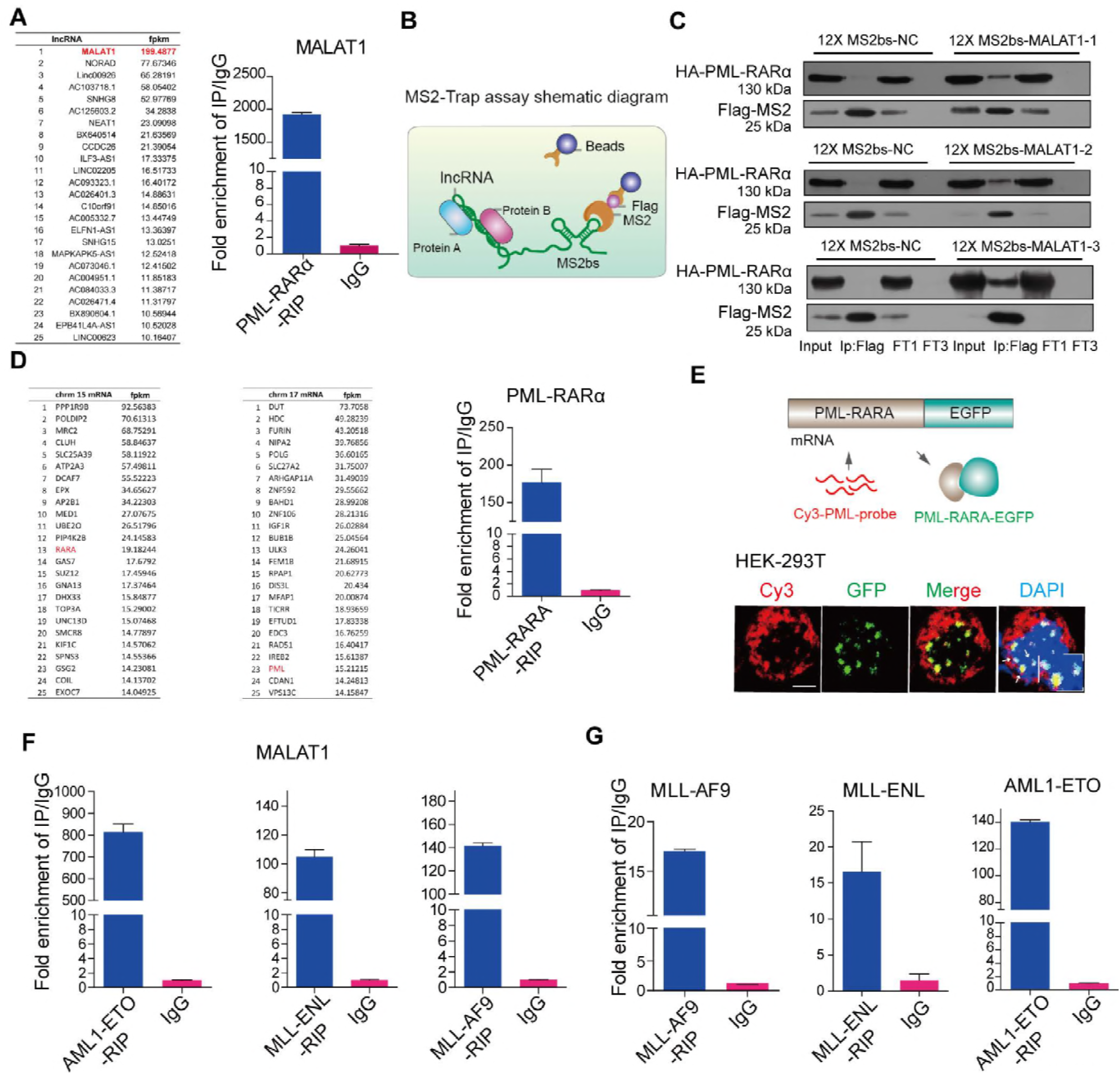
-Both the mRNA and fusion proteins were hijacked in nuclear speckles by MALAT1. **A** The mRNAs and lncRNAs binding to PML-RARα(left panel). qRT-PCR validated MALAT1 binding to PML-RARA (right panel). **B** Schematic diagram showing the procedure of MS2-Trap assays. **C** PML-RARα interacted with MALAT1 in HEK-293T cells. The three sections of MALAT1 werenamed as MALAT1–1, -2, -3 respectively (FT1, flow through; FT3, flow through after three times wash) **D** The mRNAs binding to PML-RARα. qRT-PCR confirmed the binding(right panel).. **E** The mRNA FISH assay for detecting the interaction of PML-RARα and its own mRNA in HEK-293T cells in which PML-RARα-EGFP over-expressed. PML-RARα-EGFP (green) represents the fusion protein and PML-CY3 (red) represents mRNA transcripts. Scale bar represents 4 μm. **F,G** RIP assays followed by qRT-PCR analysis for the interaction of MALAT1 and the fusion protein mRNAs with fusion proteins (MLL-ENL, AML1-ETO and MLL-AF9) in HEK-293T cells. Data are shown as the means ± s.e.m.; n = 3 independent experiments.

In addition to binding to MALAT1, the RIP-seq data and qRT-PCR also showed that PML-RARα could bind to a number of mRNAs including their own mRNAs (**Fig 1D**). In situ hybridization (FISH) assays further confirmed the direct interaction between PML-RARA and its own mRNA transcripts (**Fig 1E**). To demonstrate whether this reflects a common mechanism for chromosomal translocated genes, we further examined three other fusion proteins (MLL-ENL, MLL-AF9 and AML1-ETO). These translocations are typical oncogenic fusion genes that drive specific subtypes of leukemia(Kuntimaddi et al, 2015; Muntean & Hess, 2012; Okumura et al, 2008). The results further showed that MALAT1 and their own mRNAs were significantly enriched in the cells that overexpressed these three fusion proteins AML1-ETO, MLL-ENL and MLL-AF9, respectively (**Fig 1F and H**). These observations suggested that the MALAT1 interaction may be largely conserved in different cell types with different gene translocations. Because nuclear speckles are a subnuclear structure that is related to mRNA maturation processes, such as splicing and export, both the chimeric mRNAs and fusion proteins hijacked in nuclear speckles by MALAT1 raised the question whether MALAT1 is involved in the regulation of oncogenic fusion proteins or their mRNA maintenance or export?

### Nuclear export of chimeric mRNAs depends on the MALAT1 expression levels

To address whether MALAT1 regulates oncogenic fusion proteins or their mRNA export, we knocked down MALAT1 expression and then measured PML-RARα and MLL-AF9 protein levels, respectively. We found that both protein levels decreased sharply (**Fig 2A and B; Fig EV2A and B**); however, the mRNA levels of PML-RARα and MLL-AF9 were almost identical before and after knocking down MALAT1 (**Fig 2C and D**), raising the question that the decrease of fusion protein expression might not be related to the chimeric mRNA transcription, it might be relate to the mRNA export and translation? To this end, nuclear and cytoplasmic RNAs were extracted and used for the chimeric mRNA detection. Notably, the mRNA of fusion protein PML-RARα was significantly retained in the nucleus when MALAT1 was reduced (**Fig 2E and Fig EV2C**). And this observation was further confirmed by RNA FISH experiments as well (**Fig 2F**). Moreover, in the cells that overexpressed an artificial PML-RARα-EGFP construct, knockdown of MALAT1 resulted in significant PML-RARα mRNA retention in nucleus (**Fig 2G**), indicating that the binding of fusion proteins to their own mRNA was independent of MALAT1. These results support the hypothesis that MALAT1 promotes oncogenic fusion protein expression by facilitating chimeric mRNA export from nucleus to cytoplasm.

**Figure 2.**
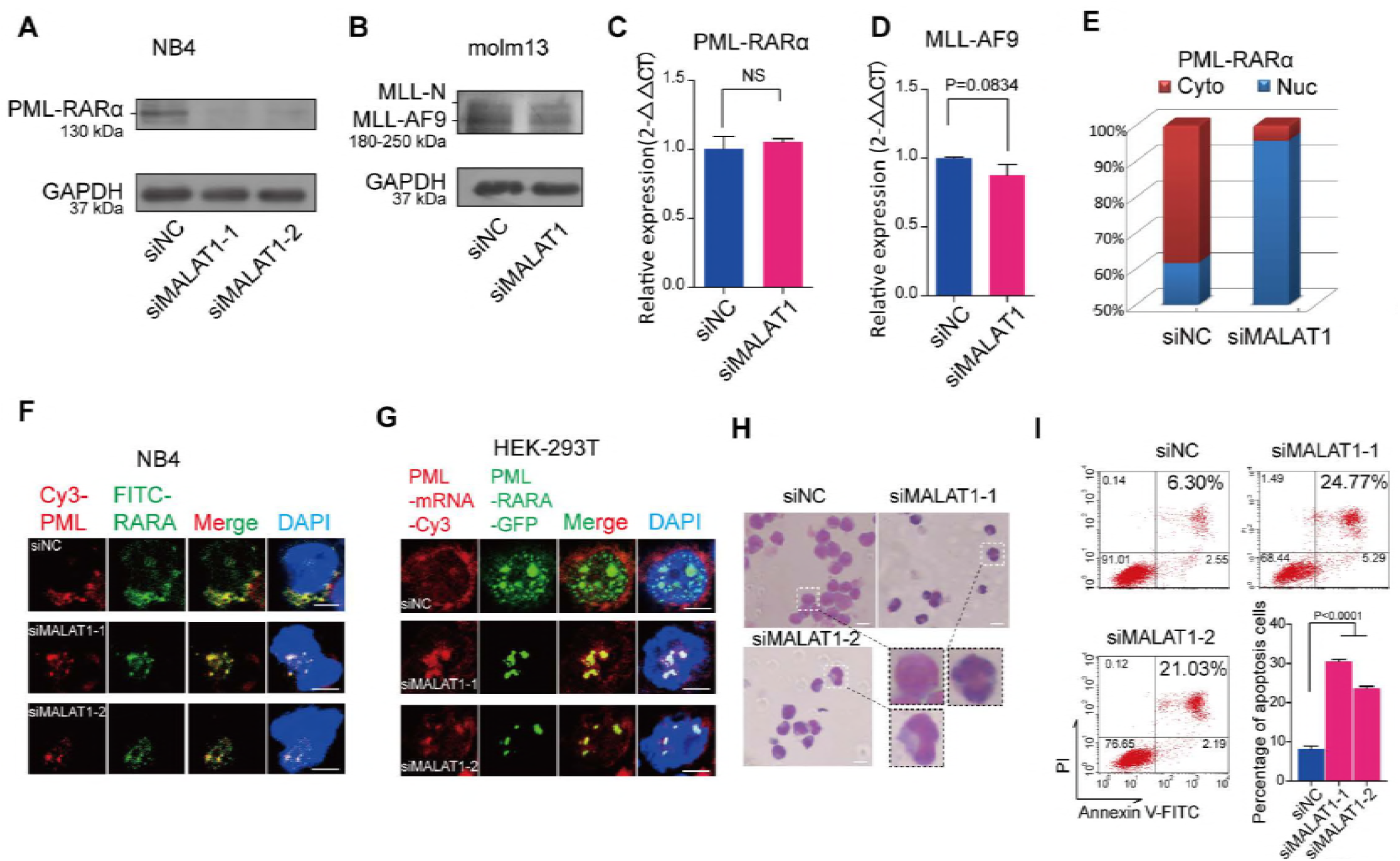
Knockdown of MALAT1 significantly repressed the expression levels of fusion proteins by promoting their mRNAs retention in nuclei. A, B Protein levels of PML-RARα and MLL-AF9 were reduced after knockdown of MALAT1 in NB4 cells and molm13 cells, respectively. GAPDH was used as a loading reference. All experiments were performed independently at least three times. C, D qRT-PCR showed the mRNA levels of PML-RARα and MLL-AF9 were not markedly changed after reducing MALAT1 levels via siRNA. Data are shown as the means ± s.e.m.; n = 3 independent experiments. **(E)** PML-RARα mRNA detection by qRT-PCR after separating the nuclear and cytoplasmic RNAs.. **F** PML-RARα mRNA location detecting by RNA FISH in NB4 cells. Two different fluorophores were added to PML (Cy3, red) and RARα (FITC, green) mRNA, respectively. The dots merged into yellow dots represent the mRNAs of PML-RARα. Scale bar represents 4 μm. **G** RNA FISH experiments were carried out according to the process shown in Figure 1b in HEK-293T cells. Scale bar represents 4 μm. **H** Wright-Giemsa staining showed that cellular differentiation was significantly promoted after reducing MALAT1 levels in NB4 cells. Scale bar represents 10 μm. **I** Cellular apoptosis measurement by flow cytometry. Data are shown as the means ± s.e.m.; n = 3 independent experiments.

It has been well documented that oncogenic fusion proteins are key drivers of oncogenesis via the blockade of myeloid cell differentiation in myeloid leukemia (Kuntimaddi et al, 2015; Muntean & Hess, 2012; Okumura et al, 2008). Therefore, we investigated whether MALAT1 also plays a role in malignant progression. Knockdown of MALAT1 prominently facilitated the differentiation of progenitor cells to monocytes and granulocytes in PML-RARα-positive cells (**Fig EV2D and E**). Karyotype analysis by Wright-Giemsa staining showed that the nuclei appeared to be significantly lobulated when MALAT1 was suppressed (**Fig 2H**) and apoptosis was significantly enhanced (**Fig 2I**). In summary, we concluded that MALAT1 maintained protein expression levels and oncogenic functions of fusion proteins by promoting their mRNA export from nucleus to cytoplasm.

### MALAT1 regulates chimeric mRNA export in a manner dependent on the YTHDC1 and SRSF3

We next asked how MALAT1 controls these chimeric mRNA export processes from nucleus to cytoplasm and regulates myeloid cell differentiation. Most mRNA export is specifically controlled by various protein factors and modifications of pre-mRNA, such as TAP-p15 heterodimers (Katahira et al, 1999; Santos-Rosa et al, 1998), adaptor proteins(Katahira, 2015) and m6A modification. Adaptor proteins include transcription-export-1 (TREX-1) complex(Chi et al, 2013; Dias et al, 2010; Reed & Cheng, 2005), TREX-2 complex(Bhatia et al, 2014), and serine-arginine rich (SR) proteins(Muller-McNicoll et al, 2016; Zhong et al, 2009). YTHDC1(Xiao et al, 2016), a ‘reader’ of m6A methylation, has been recently shown to regulate mRNA export processes by interacting with SR proteins(Roundtree et al, 2017). Therefore, we chose 13 adaptor proteins (Table EV1) and used RNAi approaches to determine which adaptor is responsible for the chimeric mRNA export process in NB4 cells(the knockdown efficiency of these mRNAs have been shown in Figure EV3A-C). Among the 13 adaptors, we found that, only in the SRSF3 and YTHDC1 knockdown cells, the amount of PML-RARα mRNA in nucleus was decreased significantly compared with that in the control (**Fig 3A and B**), suggesting that SRSF3 and YTHDC1 could be the potential adaptor proteins responsible for PML-RARα export in NB4 cells. Furthermore, the protein level of PML-RARα was sharply decreased after YTHDC1 or SRSF3 expression was reduced (**Fig 3C**). We next performed Wright-Giemsa staining assays and flow cytometry to evaluate whether the YTHDC1 and SRSF3 complex is a key regulator in cellular functions. The results showed that cellular differentiation was markedly enhanced after knocking down YTHDC1 and SRSF3 (**Fig 3D-F**), showing their regulatory roles in myeloid cell differentiation.

**Figure 3.**
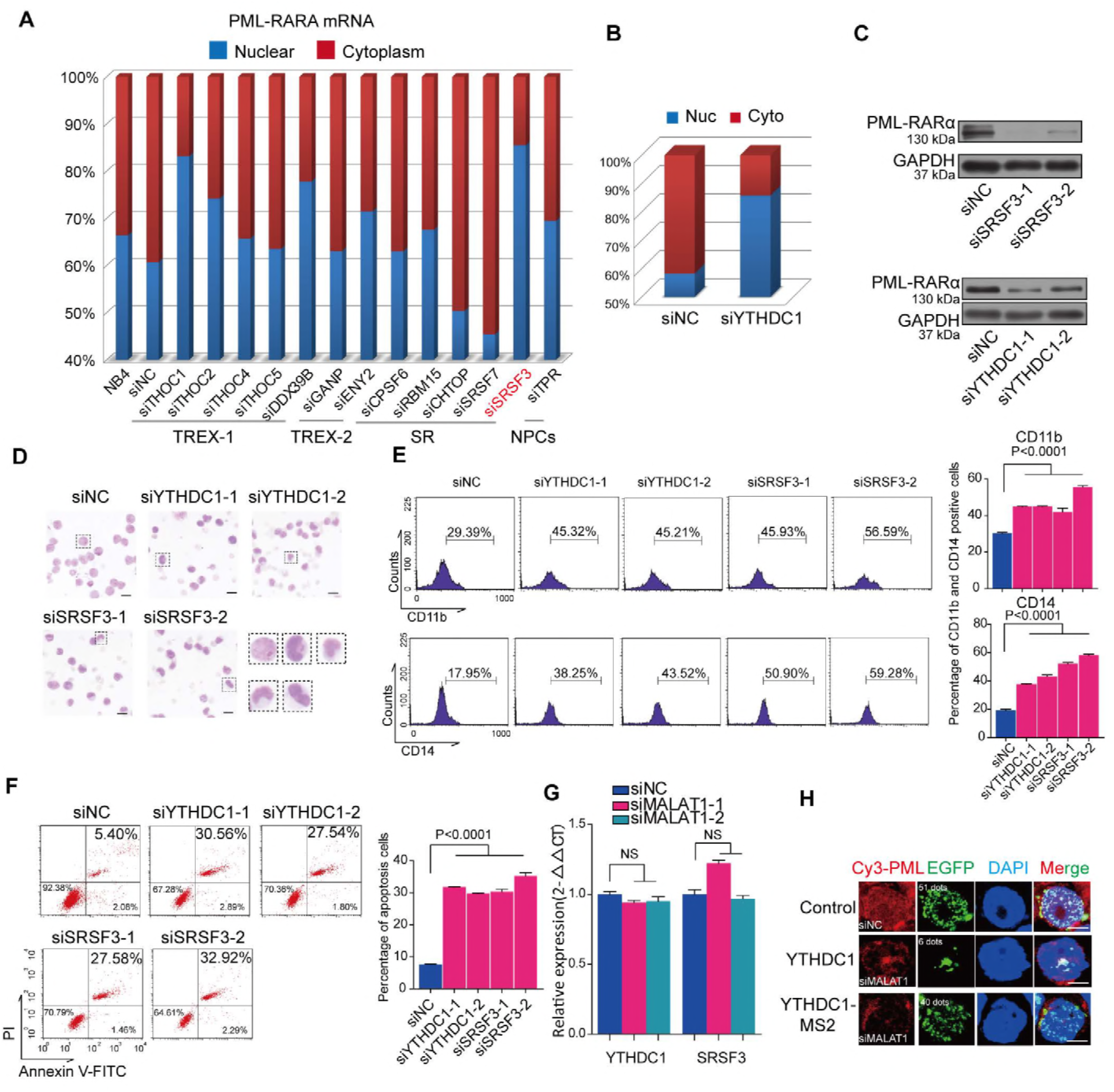
YTHDC1 and SRSF3 are responsible for the mRNA fusion gene export process. **A** Screening of mRNA export-related proteins that are specifically responsible for PML-RARα mRNA. **B** YTHDC1, a protein that regulates mRNA export by interacting with SRSF3, is related to PML-RARA mRNA export as well. **C** PML-RARα was diminished after knockdown of SRSF3 or YTHDC1 in NB4 cells. The data were obtained from three independent experiments. **D** Wright-Giemsa staining assays showed that phenotypes of the nuclei changed to a lobulated morphology after inhibiting YTHDC1 or SRSF33. Scale bar represents 10 μm. **(E)** Cellular differentiation marker (CD11b and CD14) testing by flow cytometry. **(F)** Cellular apoptosis detection by flow cytometry. **G** YTHDC1 and SRSF3 transcripts expressions do not change significantly after knockdown of MALAT1 in NB4 cells. **H** Fusion mRNA transport process was dependent on YTHDC1. RNA FISH assay for testing PML-RARα mRNA transport by transfecting constructs of PML-RARα-EGFP-12×MS2bs and YTHDC1-MS2 upon knocking down MALAT1 in HEK-293T cells. Scale bar represents 4 μm. Data are shown as the means ± s.e.m.; n = 3 independent experiments.

We next assessed the regulatory relationship between MALAT1 and YTHDC1 or SRSF3. It was shown that no significant alterations of YTHDC1 and SRSF3 levels were found following reduction of MALAT1 levels (**Fig 3G**). In addition, we designed a reporter vector to demonstrate whether the regulation of MALAT1 on the mRNA export process is dependent on the YTHDC1 recognition (**Fig 3H**). The results showed that overexpression of YTHDC1 did not change the nuclear retention of PML-RARα mRNA after knockdown of MALAT1 (**Fig 3H**), while ectopic expression of MS2-tagged YTHDC1 could significantly enhance the export of PML-RARα mRNA, illustrating that YTHDC1 is crucial for the nuclear export of mRNA transcribed by fusion genes. These results suggest that MALAT1 might regulate chimeric mRNA export and hematopoietic differentiation in a manner dependent on the YTHDC1 and SRSF3 export-associated complex, which are not regulated by MALAT1 directly.

### YTHDC1 recognition of m6A modification on PML-RARα mRNA is required for its export

YTHDC1, a member of the YTH domain-containing protein family, is reported to be an important m6A reader. Interestingly, a recent study showed that YTHDC1 facilitated m6A-methylated mRNA export by interacting with SRSF3; therefore, we proposed that the PML-RARα mRNA export process might be regulated by m6A modification. To evaluate this hypothesis, we first analyzed the sequence of PML-RARα mRNA to determine the conserved m6A motif. A few potential motifs for METTL14 and WTAP in the junction sites of PML-RARα (exons 4, 5 and 6 of the PML gene even including introns) were found (**Fig 4A** and **Table EV2**). Then, we performed the MeRIP assay with or without knockdown of MALAT1 and m6A methyltransferases, respectively. PCR results showed that the band representing sequences containing one of the motifs of METTL14 in PML exons 4, ‘GGACC’, was diminished following the reduction in m6A levels by knocking down methyltransferases (**Fig 4A and Fig EV4A**), indicating that the ‘A’ base in the motif was indeed modified by m6A methyltransferases (**Fig EV4B**). Furthermore, the PML exon 4 m6A signal was diminished in the MALAT1-knockdown samples as well (**Fig 4A and B**).

**Fig. 4.**
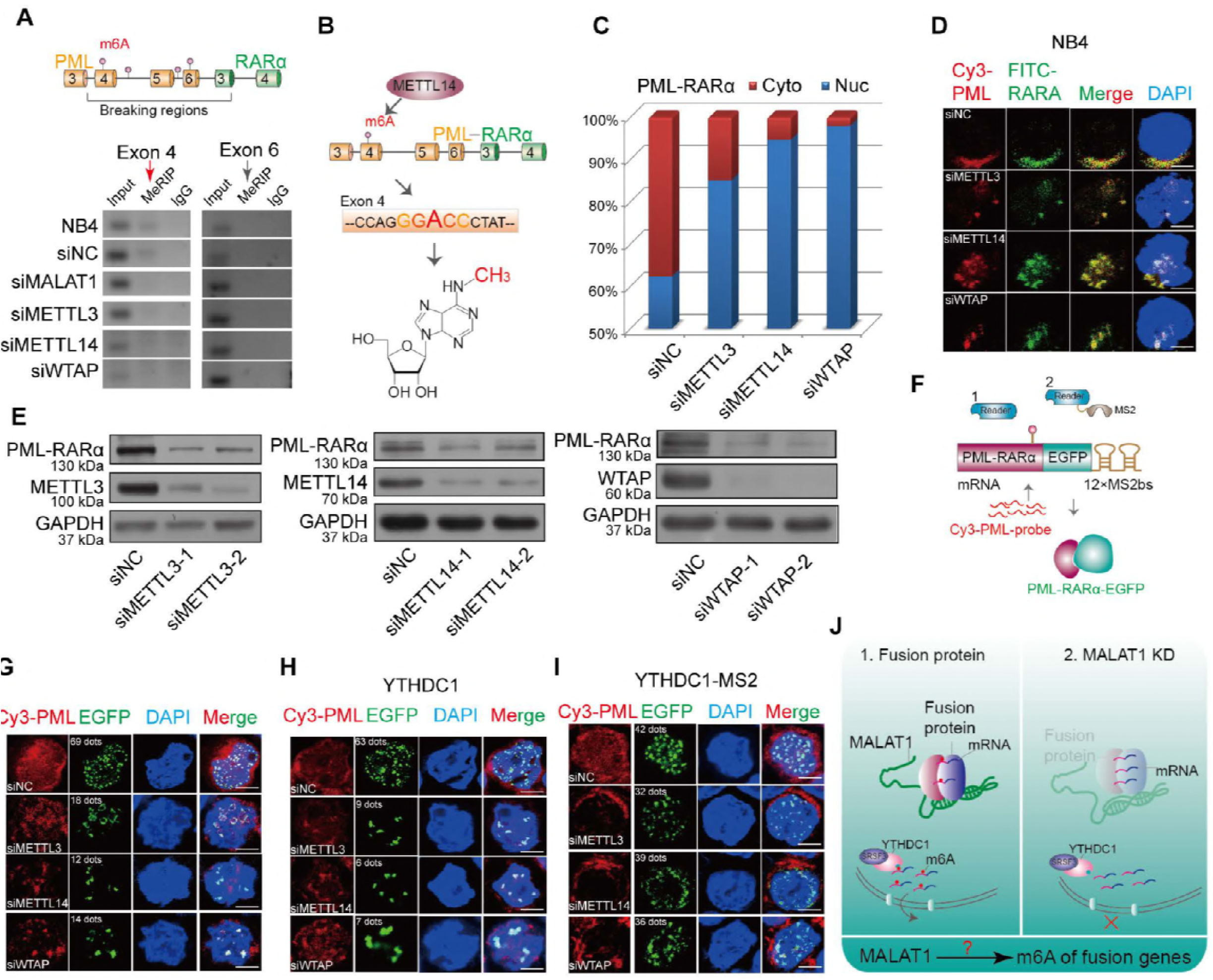
MALAT1-regulated m6A modification in the nucleus is responsible for the fusion gene export by YTHDC1. **A** A schematic showing potential m6A motifs in regions of PML-RARα mRNA. MeRIP assays were performed in NB4 cells after knockdown of m6A methyltransferases, MALAT1 or a negative control with siRNAs. Then, PCR sequencing for these sites, including potential motifs, were carried out; agarose electrophoresis was shown. **B** Schematic diagram of METTL14-methylated m6A sites in PML-RARα mRNA exon 4. **C** PML-RARA mRNA dispersion in nuclei and cytoplasm was detected after reducing the m6A level by knocking down the methyltransferases. **D** RNA FISH analysis showing the PML-RARα mRNA location in cells by two probes, Cy3-PML (red) and FITC-RARα (green). Scale bar represents 4 μm. **E** The protein levels of PML-RARα, METTL3, METTL14 and WTAP were monitored after knockdown of m6A methyltransferases by several different siRNAs. Three independent experiments were carried out. **F** Schematic diagram showing RNA FISH assays for testing PML-RARα mRNA transport by transfecting PML-RARα-EGFP-12×MS2bs and YTHDC1-MS2 constructs with knockdown of m6A in HEK-293T cells **G** Representative images showed fusion protein green dots were reduced. Red dots were remained in nucleus when m6A methyltransferases were knocked down in HEK-293T cells. Scale bar represents 4 μm. **H** Combined with cotransfecting YTHDC1 into HEK-293T cells, representative images of PML-RARα mRNA and PML-RARα-EGFP are shown in HEK-293T cells after reducing m6A levels. Scale bar represents 4 μm. **I** Combined with cotransfecting YTHDC1-MS2, representative images showed PML-RARα mRNA retention in nuclei was significantly restored after reducing m6A, as compared with the negative controls. Scale bar represents 4 μm. **J** Schematic diagram showing the hypothesis the potential roles of MALAT1, m6A and the reader YTHDC1 in the fusion gene export process.

Consistently, as shown in **Fig 4C** and **Fig EV4C and D**, the diminished of m6A modification retained the PML-RARα in nucleus, and the total mRNA levels were unchanged. Similar results were obtained suing RNA FISH experiments upon m6A methyltransferases or MALAT1 knockdown (**Fig 4D and Fig 2F**). Moreover, both PML-RARα and MLL-AF9 protein levels were reduced significantly after reducing the expression of m6A ‘writers’ (**Fig 4E and Fig EV4F**). M6A knockdown significantly increased nuclear PML-RARα mRNA levels in HEK-293T cells that overexpressed the PML-RARα construct (**Fig 4G**). Furthermore, overexpressed YTHDC1 did not change the nuclear retention of PML-RARα mRNA after knockdown of m6A complex genes METTL3, METTL14 as well as WTAP (**Fig 4H**), whereas forced expression of YTHDC1 (MS2-tagged) could significantly restore the levels of PML-RARα mRNA export; this finding indicated that YTHDC1 regulates nuclear export of mRNA of fusion genes in a m6A-dependent manner (**Fig 4I**). Additionally, Wright-Giemsa staining assays showed that nuclei appeared to be significantly lobulated when MALAT1-regulated m6A methyltransferases were suppressed (**Fig EV4G**).

In conclusion, m6A modification of PML-RARα mRNA, controlled by MALAT1, is a key signature for PML-RARα mRNA processing in nuclear speckles. MALAT1-regulated m6A modification, which is recognized by YTHDC1, is necessary for the export of PML-RARα and cellular functionality (**Fig 4J**).

### MALAT1 and fusion protein complexes serve as a functional loading bridge for the interaction of chimeric mRNA and METTL14

The above results demonstrated that chimeric mRNA export is modulated by MALAT1 and m6A modification, which is recognized by YTHDC1. We next asked how MALAT1 regulates the m6A modification levels of fusion transcripts. Does MALAT1 affect the m6A methyltransferase expression levels? So we assessed m6A methyltransferase (METTL3, METTL14 and WTAP) levels following a reduction in MALAT1 expression in NB4 cells. However, the expression levels of the three m6A methyltransferases were not altered (**Fig 5A**), and dot blot assays further supported this observation (**Fig EV5A**), indicating that m6A modification levels are not altered by MALAT1 directly. It has been previously reported that MALAT1 could bind to one of the m6A RNA methyltransferase members, WTAP(Horiuchi et al, 2013). Importantly, m6A methyltransferases are also located in nuclear speckles(Liu et al, 2014). Thus, we hypothesized that MALAT1 might hijack fusion proteins in nuclear speckles to interact with the m6A modification complex and, in turn, regulate PML-RARα mRNA modification and export. Thus, we tested the interaction of m6A methyltransferases with MALAT1, as well as with fusion proteins. MS2-Trap assays showed that the m6A methyltransferases METTLE 3 and 14 could directly bind to three sections of MALAT1 (**Fig 5B and Fig EV5B**). With immunofluorescent labeling assays in NB4 cells and co-immunoprecipitation assays in HEK-293T cells, we found that PML-RARα and m6A methyltransferases also form a complex (**Fig 5C and D**). We also constructed lentivirus-infected NB4 cells that stably express flag-tagged METTL3, METTL14 and WTAP to further validate this interplay. **Fig EV5C** showed the binding of flag-tagged m6A methyltransferases and endogenous PML-RARα, which is consistent with that of antibodies for endogenous METTL3 and WTAP proteins.

**Figure 5.**
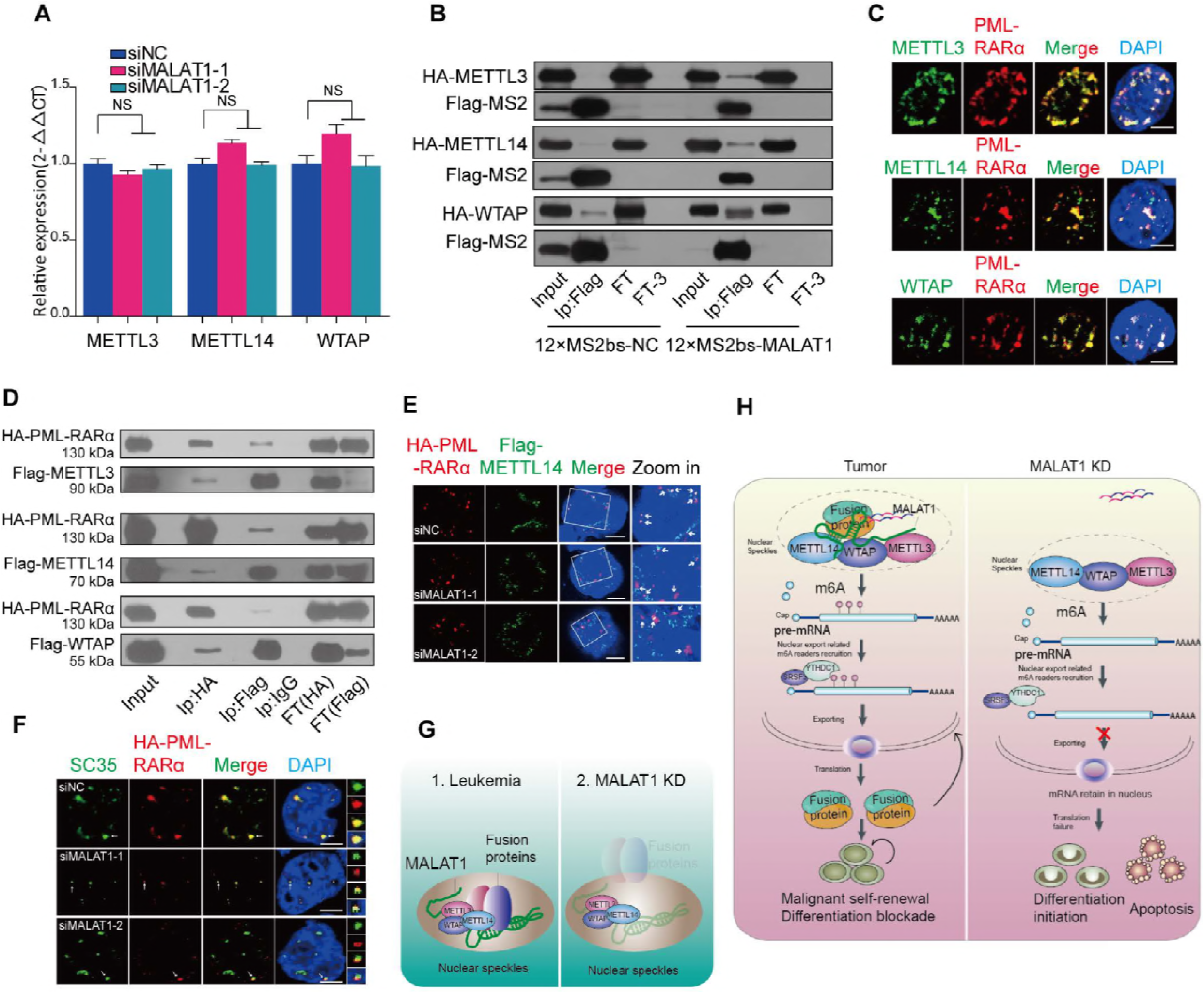
MALAT1 forms a complex with fusion proteins and m6A methyltransferases. **A** m6A methyltransferase expression was not regulated by MALAT1. Data are shown as the means ± s.e.m.; n = 3 independent experiments. **B** MALAT1 bound to m6A methyltransferases METTL3, 14 and WTAP directly by MS2-Trap assays in HEK-293T cells(FT1, flow through; FT3, flow through after three times wash).The experiments were performed independently at least three times. **C** Immunofluorescence assays by laser confocal microscopy showed PML-RARα (anti-RARα) and m6A methyltransferases (anti-METTL3, 14 and WTAP) co-localized in NB4 cells. Scale bar represents 4 μm. **D** Co-immunoprecipitation and Western blot experiments were used to further detect the interaction of PML-RARα and m6A methyltransferases in HEK-293T cells(FT1, flow through; FT3, flow through after three times wash). **E** HA-PML-RARα (anti-HA) and Flag-METTL14 (anti-Flag) localized in different positions in nuclei following knockdown of MALAT1 in HEK-293T cells by the methods of immunofluorescence. Scale bar represents 4 μm. **F** The nuclear localization of HA-PML-RARα (anti-HA) and nuclear speckle marker SC35 (anti-SC35) was only partially overlapping in the siMALAT1 samples compared to the negative controls in immunofluorescence assays. Representative images from confocal microscopy are shown from three independent experiments, scale bar represents 4 μm. **G** Schematic models indicating the potential mechanistic roles of fusion proteins and m6A methyltransferases in nuclear speckles. **H** Schematic models showing the mechanism of MALAT1-mediated regulation of the fusion gene export process by modulating the interaction of fusion proteins with METTL3, 14 and WTAP during hematopoiesis.

Next, we investigated if MALAT1 mediates the interaction between PML-RARα and m6A complex components. The co-immunoprecipitation assays showed that the binding of METTL14 to PML-RARα was significantly reduced when MALAT1 level was decreased; however, the binding to METTL3 and WTAP was not significantly affected (**Fig EV5D**), indicating that METT14 is the main m6A methyltransferase component that might be of importance for PML-RARα mRNA modification, which is consistent with the data shown in **Fig 4B** that METT14-modified motif sequence was found in PML-RARA mRNA. Immunofluorescence assays also showed that the overlapping areas of METTL14 and PML-RARα were reduced after treatment with siRNAs targeting MALAT1 (**Fig 5E**). Very interestingly, when MALAT1 expression levels were reduced, PML-RARα dispersed near the nuclear speckles as a partially localized structure in HEK-293T cells with ectopic expression of PML-RARα (**Fig 5F**), suggesting that MALAT1 might maintain their localization in nuclear speckles and their interaction with METTL14 (**Fig 5G**). Therefore, MALAT1 and fusion protein complexes may serve as a functional loading bridge for the interaction of PML-RARα mRNA and m6A methyltransferases, further regulating modification and PML-RARα mRNA export (**Fig 5H**).

### MALAT1 reduced fusion protein expression levels in primary cells from patients and extended survival in a mouse model

Finally, we used primary cells from patients with different gene translocations for functional validation. We collected samples from clinical patients with PML-RARα. Upon knockdown of either MALAT1 or METTL3, the levels of PML-RARαwere decreased significantly (**Fig 6A**). We further established an ascites NOD-SCID mouse model using human leukemia cell line NB4 harboring PML-RARα to reveal the function of MALAT1 and m6A in vivo (**Fig 6B**). Ascites NOD-SCID mice were produced 20 days after injecting 5 × 10^6^ lentivirus-infected NB4 cells with specific knockdown of MALAT1 (named NB4-Lv-NC or NB4-Lv-shMALAT1) or 5 × 10^6^ lentivirus-infected NB4 cells that overexpressed m6A methyltransferases (METTL3, METTL14 and WTAP), respectively. The stable knockdown efficiencies of NB4-Lv-shMALAT1 and NB4-Lv-METT3, METT14 and WATP were shown in **Fig EV 5C, 6A, B**. The ascites fluid was collected and used for Western blot and FACS analysis and Wright–Giemsa staining. Western blot analysis showed that PML-RARα was reduced when MALAT1 was knocked down in ascites fluid of mice (shMALAT1-M) (**Fig 6C and Fig EV6C**), whereas its level was markedly increased in the ascites fluid of m6A-enhanced NOD-SCID mice compared to that of controls (**Fig 6D**). The nuclei were found to be lobulated and differentiated by Wright–Giemsa staining (**Fig EV6D and E**). FACS analysis showed that CD11b and CD14 expression level were significantly induced (**Fig 6E and F; Fig EV6F**). To further determine the effects of MALAT1 on PML-RARα in vivo, we also injected a NOD-SCID xenograft mice through tail vein with NB4 cells and moml13 cells that harboring MALAT1 short hairpin RNA (shMALAT1) or control short hairpin RNA (shNC). The mice were sacrificed after 21 days and the percentages of GFP^+^ cells were decreased in the bone marrow (BM) of mice treated with either the shMALAT1 NB4 cells or the shMALAT1 molm13 cells compared with those in control mice ( **Fig 6G and Fig EV6G, lef**t). In line with the CD11b expressions measured by flow-cytometry, reducing the level of MALAT1 significantly promote cell differentiation in the murine model, and another marker CD14 for differentiated cells confirms this phenomenon as well (**Fig 6G and Fig EV6G, right**). Consistent with the pro-differentiation phenotype in vitro, impaired expression levels of the fusion protein PML-RARα were observed in BM cells of NB4-Lv-shMALAT1 injected mice rather than fusion protein levels in the BM cells of control mice (**Fig 6H**) Wright-Giemsa staining also verified that human leukemic cell, which are larger than bone marrow cells of mice, were induced to differentiate upon knocking down MALAT1 (**Fig 6I and Fig EV6H**). MALAT1 knockdown mice had extended survival than did the control mice (**Fig 6I**).

**Figure 6.**
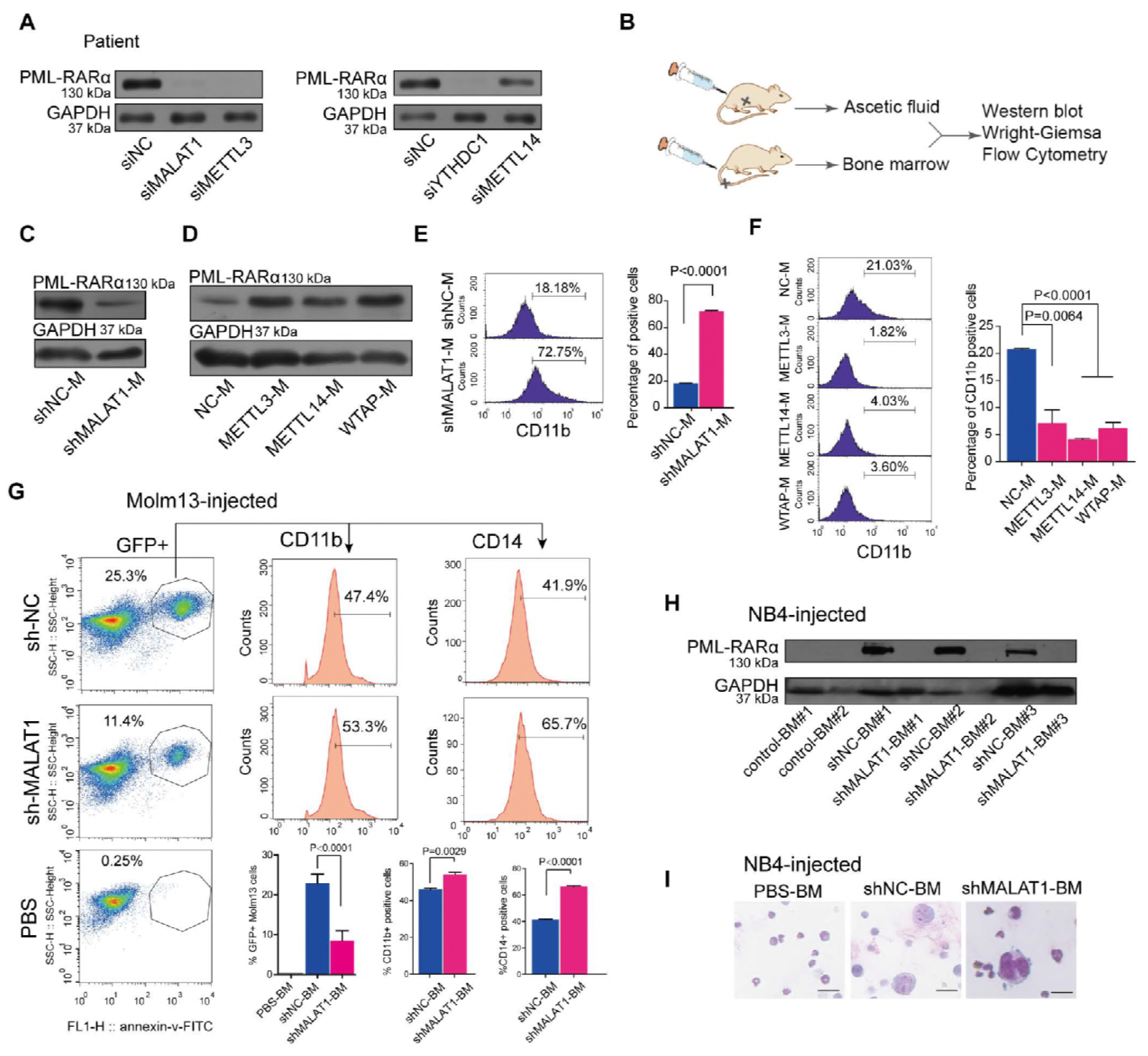
MALAT1 functions in cellular differentiation via m6A during malignant hematopoiesis in vivo. **(A** The protein levels of PML-RARA in clinical patient #1 were determined via Western blot after knocking down MALAT1, m6A methyltransferases and YTHDC1 with siRNAs. **B** Schematic diagram showing NOD-SCID murine models of ascites fluid and tail vein injection strategy, followed by Western blot, Wright-Giemsa and flow cytometry analysis. **CD** Western blot measurements of PML-RARA protein levels in the ascites fluid of mice injected with NB4 cells with suppressed MALAT1 (shMALAT1-M) or overexpressed m6A methyltransferases (METTL3-M, METTL14-M, WTAP-M). Three independent experiments were performed. **E,F** The differentiation marker CD11b of NOD-SCID murine ascites fluid was analyzed by flow cytometry after altering MALAT1 expression or m6A levels. Data are shown as the means ± s.e.m.; n = 3 independent experiments. **G** Differentiation marker CD11b and CD14 of BM cells of NOD-SCID mice intravenously injected by MLL-AF9 positive molm13 cells were analyzed by flow cytometery after reducing MALAT1.3 mice each group, the mice injected the PBS as the negative control (n=1). Data are analyzed and shown as the mean ± s.e.m.; n = 3 independent experiments. **H** In the tail vein injection mouse model, bone marrow cells were used to detect PML-RARA protein levels by Western blot after suppressing MALAT1 expression (shMALAT1-BM#1,2,3). **I** Wright-Giemsa assays were carried out to verify that human NB4 cells (shNC and shMALAT1), which were much larger than mouse bone marrow cells, were transplanted to bone marrow. IScale bar represents 10 μm.

Collectively, we established that a functional RNP complex, which composed of MALAT1, leukemia-related fusion proteins and m6A methyltransferases, regulates the mRNA transcript export process in nuclear speckles. We revealed a lncRNA-fusion protein-m6A autoregulatory loop controls the chimeric mRNA export in malignant hematopoiesis (as shown in **Fig 5 H**).

## Discussion

Fusion proteins resulting from aberrant chromosomal translocations have been determined to be key drivers for tumorigenesis(Krivtsov & Armstrong, 2007; Mertens et al, 2015; Mitelman et al, 2007). In this study, it is the first to report that these fusion proteins were found to be located in nuclear speckles and physically interact with m6A methyltransferases (e.g., METTL3, METTL14 and WTAP) through the lncRNA MALAT1, an important regulatory component of the RNA-protein complex in nuclear speckles. We further discovered that the chimeric mRNA export was regulated by MALAT1 via m6A modification on their transcripts. Silencing MALAT1 could affect the export of these chimeric mRNAs from nucleus to cytoplasm, inhibiting the translation of fusion genes and ultimately controlling cell fates. The findings in this study showed that MALAT1 may be a potential therapeutic target for the treatment of fusion protein-driven cancers. Furthermore, the study is the first to show lncRNA regulation of the chimeric mRNA export process.

LncRNAs, a subclass of ncRNAs with a sequence length greater than 200 nt, have been shown to have multiple functions in various cellular processes through binding to chromatin regulatory proteins(Bhatia et al, 2014; Brown et al, 2016), DNA and even RNA on the basis of their secondary structures and/or primary sequences(Okumura et al, 2008). LncRNAs are also responsible for retaining of double-stranded mRNAs in nucleus(Chen & Carmichael, 2009; Edupuganti et al, 2017; Fox & Lamond, 2010; Scadden, 2009). For instance, NEAT1, a constitutive lncRNA in the nuclear paraspeckle, is responsible for the nuclear retention of inverted-repeat Alu element (IRAlu, double-stranded RNA structure) mRNAs by interacting with the A-I editing protein P54NRB. In contrast to the previously identified functions in the regulation of the alternative splicing process in nuclear speckles, MALAT1 is shown here to play a key role in the mRNA export process for the first time. We found that MALAT1regulates typical chimeric mRNA export and determines lineage differentiation by directly binding to a complex of RNA modification-related proteins responsible for retaining double-stranded mRNAs in the nucleus.

Previous studies have shown that m6A modifications on the lncRNA XIST promote XIST-mediated X chromosomal transcriptional repression by interacting with the RNA binding proteins RBM15 and RBM15B(Chen & Carmichael, 2009; Edupuganti et al, 2017; Fox & Lamond, 2010; Scadden, 2009), as well as the m6A methyltransferase WTAP(Horiuchi et al, 2013). Here, we found that MALAT1 induces specific m6A modification on the mRNAs by binding to these specific fusion proteins and m6A methyltransferases. Interestingly, it has been reported that a m6A modification at the MALAT1-2577 site, functioning as an RNA structural switch, regulates RNA-protein interactions(Brown et al, 2014; Brown et al, 2016). Whether this m6A site contributes to the MALAT1-induced RNA and protein complex formation and function needs to be further investigated.

Finally, the study revealed the mechanism of the mRNA export process of chimeric mRNA includes lncRNA interactions with oncogenic fusion proteins, which in return autoregulate the m6A levels of their own mRNAs. The accurate intracellular interaction of proteins and their own mRNA has recently become regarded as a pervasive yet significant molecular mechanism(Terenzio et al, 2018; Wu et al, 2015). More and more studies have shown that this interaction is a functional and regulatory model. Similarly, our work here revealed that the fusion protein localization and interaction with other regulatory proteins in nuclear speckles enables their own mRNA export from the nucleus to the cytoplasm. Furthermore, we found the lncRNA MALAT1 determined the interaction with RNA methyltransferases, resulting in the control of m6A levels in chimeric mRNAs that bound to proteins. Similar to the regulation of protein interactions by lncRNAs, overexpression of another ncRNA, e.g., pre-microRNA, resulted in nuclear mRNA retention and subsequent protein reduction of human Dicer levels (required for miRNA maturation) via competitive binding to Exportin-5 with Dicer mRNA(Yi et al, 2003). In this study, we revealed that fusion proteins interact with RNA methyltransferases and subsequently autoregulate the export of their own mRNAs, binding to them via m6A. It could be concluded that, in the malignant hematopoiesis process, translocated proteins appear to form a positive feedback loop to maintain their persistent and ectopic expression by binding to their own mRNAs and promoting transport epigenetically. This study revealed that the novel lncRNA-fusion protein-m6A autoregulatory loop systems contribute to the chimeric mRNA export process and their eventual malignant functions.

In summary, this study showed that the nuclear lncRNA MALAT1 regulates the mRNA transport process from the nucleus to the cytoplasm via m6A modifications. MALAT1 promotes PML-RARA, MLL-ENL, MLL-AF9 and other fusion proteins to function in a dimensional RNA layer and further regulate the mRNA export process via the potential utility of chemical modifications of the mRNA. MALAT1, acting as a platform, modulates different translocations, resulting in oncogenic protein colocalization with m6A methyltransferases in nuclear speckles. YTHDC1, an m6A reader, regulates these chimeric mRNA export processes and functions in fusion protein-driven differentiation blockade and apoptosis inhibition. Targeting MALAT1 or the m6A reader YTHDC1 could retain oncogenic gene mRNAs in nuclei and contribute to the subsequent elimination of abnormally mutated malignant cells. The pathway described here is critical for leukemia and reveals a novel mechanism of lncRNA-mediated m6A modification and mRNA export of chimeric mRNA, shedding light on the multifunctional regulatory roles of lncRNAs in carcinogenesis. Therefore, targeting MALAT1 will be a promising therapeutic target for inhibiting fusion protein-driven cancer progression.

## Materials and Methods

### Cell lines and cultures

Leukemia cells (NB4, Molm13, Kasumi-1 and K562) and HEK-293T cells were purchased from American Type Culture Collection (ATCC, USA.) and were cultured in RPMI Medium Modified (HyClone, USA) and DMEM (HyClone, USA), respectively, containing 10-20% fetal bovine serum (Gibco, Thermo Fisher Scientific,USA). The cells were cultured in a humidified atmosphere containing 5% CO2 at 37°C. Puromycin was dissolved in DMSO ultrapure water with the concentrations of 10 mg/ml. Wright-Giesma staining was used to test for changes in cell morphology.

### FACS analysis

Cell differentiation was assessed by detecting the surface antigen expression of CD11b and CD14 (eBioscience, USA) by flow cytometric analysis using a BD FACSAria cytometer (BD Biosciences, USA).

### RNA isolation and quantitative real-time PCR

Total RNA from leukemia cells was extracted using TRIzol reagent (Invitrogen, USA) according to the manufacturer’s guidelines. RNA separation of nucleus and cytoplasm is performed according to the previous report methods(Horiuchi et al, 2013). Real-time PCR was performed to quantify mRNA expression using SYBR^®^ Premix Ex TaqTM (TliRNaseH Plus) (Takara, Japan) according to the manufacturer’s instructions. The qRT-PCR was performed to detect different fusion protein transcripts, m6A methyltransferases mRNA and lncRNA MALAT1 expressions. Briefly, RNA was reverse-transcribed to cDNA using a PrimeScriptTM RT reagent kit with gDNA Eraser (Perfect Real Time) (Takara, Japan) and amplified with specific mRNA RT and PCR amplification primers. GAPDH served as internal controls. For the fusion proteins’ mRNAs, such as *PML-RARα, MLL-AF9, BCR-ABL1 and AML1-ETO*, primers were designed as previously reported. The oligonucleotide sequences are shown in **Supplementary Table 3**.

### Constructs

The human METTL3, METTL14 and WTAP CDS were obtained in human cells and the primers were designed and synthesized according to the previous reports. PML-RARA full length was amplified based on the total RNA from the NB4 cells. MLL-ENL, AML1-ETO, BCR-ABL full length expression plasmid were purchased from addgene. All the primers used for plasmid constructions were showed in Supplementary Table 3. The lentiviral shRNA expression plasmids targeting MALAT1 was performed according to the manufacture’s protocol of pGreenPuro™ shRNA Cloning and Expression Lentivector (System Biosciences, SBI). The lentiviral overexpression plasmids of METTL3, METTL14 and WTAP were designed and carried out with the PCDH lentivirus plasmid from addgene. PML-RARA-GFP gene was generated by homologous recombination and transferred to pcDNA3.1 which was also purchased from addgene. YTHDC1 full length of CDS was amplified from human NB4 cells. PCR products were obtained and then cloned into pEASY-T3 (TransGen Biotech, China) for further sequencing.

### Transient transfection

For the IF and IP assays of target proteins overexpression, HEK-293T cells were seeded in 6-well sterile plastic culture plates or a 60 mm diameter dish at a density of 3×10^5^ cells and 1.5×10^6^ per well respectively with complete growth medium. The cells were transfected with plasmids using Lipofectamine 3000 (Invitrogen) according to the manufacturer’s protocol. Small interfering RNAs (siRNAs) against human MALAT1 transcripts according to previous research and the negative control RNA duplex (RiboBio, China) were purchased from Guangzhou Ribo-Bio Co., Ltd. The siMALAT1-1 sequences were designed and expressed as shRNAs: sh-MALAT11. The siRNAs against human mRNA export adaptors were designed according to published data shown in **Supplementary Table 4**. Cells were transfected using the Neon^®^ Transfection System (Invitrogen, USA) with 100 pmol of oligonucleotides in 10 μl reactions according to the manufacturer’s guidelines.

### Western blot

Protein extracts were boiled in RIPA buffer (P0013B, Beyotime) and separated in a sodium dodecyl sulfate polyacrylamide electrophoresis (SDS-PAGE) gel. The proteins were then transferred to a polyvinylidene fluoride membrane (Millipore, USA) and probed with the primary antibodies, which are shown in **Supplementary Table 5**. Finally, the second antibody used and the reactive bands were detected by ECL (Thermo Scientific Pierce) in a dark room.

### RNA FISH

Different fluorescence labeled RNA probe targeting PML and RARα mRNAs was synthesized by Ribo-Bio (Cy3-PML mRNA) and Sangon Biotech (FITC-RARα mRNA). The targeting sequence of RARA is: ‘GCAAATACACTACGAACAACAGCTCAGAACAACGTGTC’. RNA FISH was carried out according to guidelines of Fluorescent In Situ Hybridization Kit (Ribo-Bio, China).

### Co-immunoprecipitation and MS2 Trap assay

Co-immunoprecipitation assay was performed as the published articles.

MS2 Trap procedure was carried out as previously described(Yoon et al, 2012).

### Immunofluorescence

The experiments are performed as our previous described procedures.39 EM images were obtained from thin sections using a JEM1400 electron microscope (JEOL, Akishima, Tokyo, Japan). The relative cytosolic areas in cross-sections of 10 cells were analyzed by Image-Pro Plus software. And fluorescence signals obtained using anti-FLAG (mouse) (A8592, Sigma), anti-FLAG (rabbit) (20543-1-AP, Proteintech), anti-HA (mouse) (H3663, Sigma) and anti-HA (rabbit) (C29F4, Cellular signaling), anti-SC35 antibody (Ab11826, Abcam) and anti-SF2 antibody (sc-33652, Santa Cruz) were acquired and analyzed by Carl Zeiss LSM 880 microscope (Carl Zeiss, Germany). At least 10 cells from each group were analyzed.

### RNA immunoprecipitation and sequencing

RNA immunoprecipitation was performed using a Magna RIP RNA-Binding Protein Immunoprecipitation Kit (Millipore, USA) according to the manufacturer’s instructions. Enriched total RNA was then prepared for the further sequencing and analysis of lncRNA and mRNA in Annoroad Company.

### MeRIP assay

Total RNA was isolated and then mRNA collected according to the GenElute mRNA miniprep kit (Sigma, Cat. No.MRN10). mRNA of the samples was further performed according to the protocol of Magna MeRIP™ m6A Kit Transcriptome-wide Profiling of N6-Methyladenosine (Catalog No. 17-10499)

### Animal models

Five-week-old NOD-SCID mice were maintained under specific pathogen-free conditions in the Laboratory Animal Center of Sun Yat-sen University, and all experiments were performed according to our Institutional Animal Guidelines. The human APL-ascites NOD-SCID mice model and tail vein injection model were used as previously described. The tumor xenografts were established by a single intra peritoneal transplantation of 5 × 10^6^ NB4 cells in 0.2 ml of PBS into NOD-SCID mice. The NOD-SCID mice were intravenously (tail vein) implanted with sh-RNA-established NB4 cells. Direct injection of 5×10^6^ shRNA-transformed NB4 cells in 150 μL of PBS was performed to establish intravenous (tail vein) leukemia. The xenografted mice were randomized into different groups. For the control, 150 μL of PBS without cells was injected. Subsequently, the BM from xenograft mice were treated with a red blood cell lysis buffer (Biolegend, USA), and cells washed with PBS containing 2% FBS before cell cytometry. Flow cytometry for the GFP+ *%* of transduced NB4 cells was performed on a C6 cytometer (BD, USA) and analyzed using FlowJo software. Three remaining mice were used to perform the survival assay.

### Statistical analysis

Data are expressed as the means ± SD of three independent experiments. One-way ANOVA was performed to compare multiple groups, and Tukey’s Multiple Comparison Test was used to analyze multiple comparisons. Leukemia-free survival was analyzed using the Kaplan-Meier method with a log-rank test. Two-tailed tests were used for univariate comparisons. For univariate and multivariate analysis of prognostic factors, a Cox proportional hazard regression model was used. p<0.05 was considered statistically significant.

### Data availability

The relevant data are available from the corresponding author upon reasonable request.

## Acknowledgments

This research was supported by National Key R&D Program of China (No. 2017YFA0504400) and National Natural Science Foundation of China (No. 81770174, 31700719 and 31870818).

## Author Contributions

Z. H.C designed and performed the research, analyzed data and wrote the manuscript. Z.C.Z.,T.Q. C., C. H., Y.M.S., W.H., L.Y. S., K. F. performed the research and analyzed data. X.Q.L. collected and analyzed clinical data. W.T.W. and Y.Q.C. designed the research and wrote the manuscript.

## Expanded View Figure legends

**Figure EV1.**
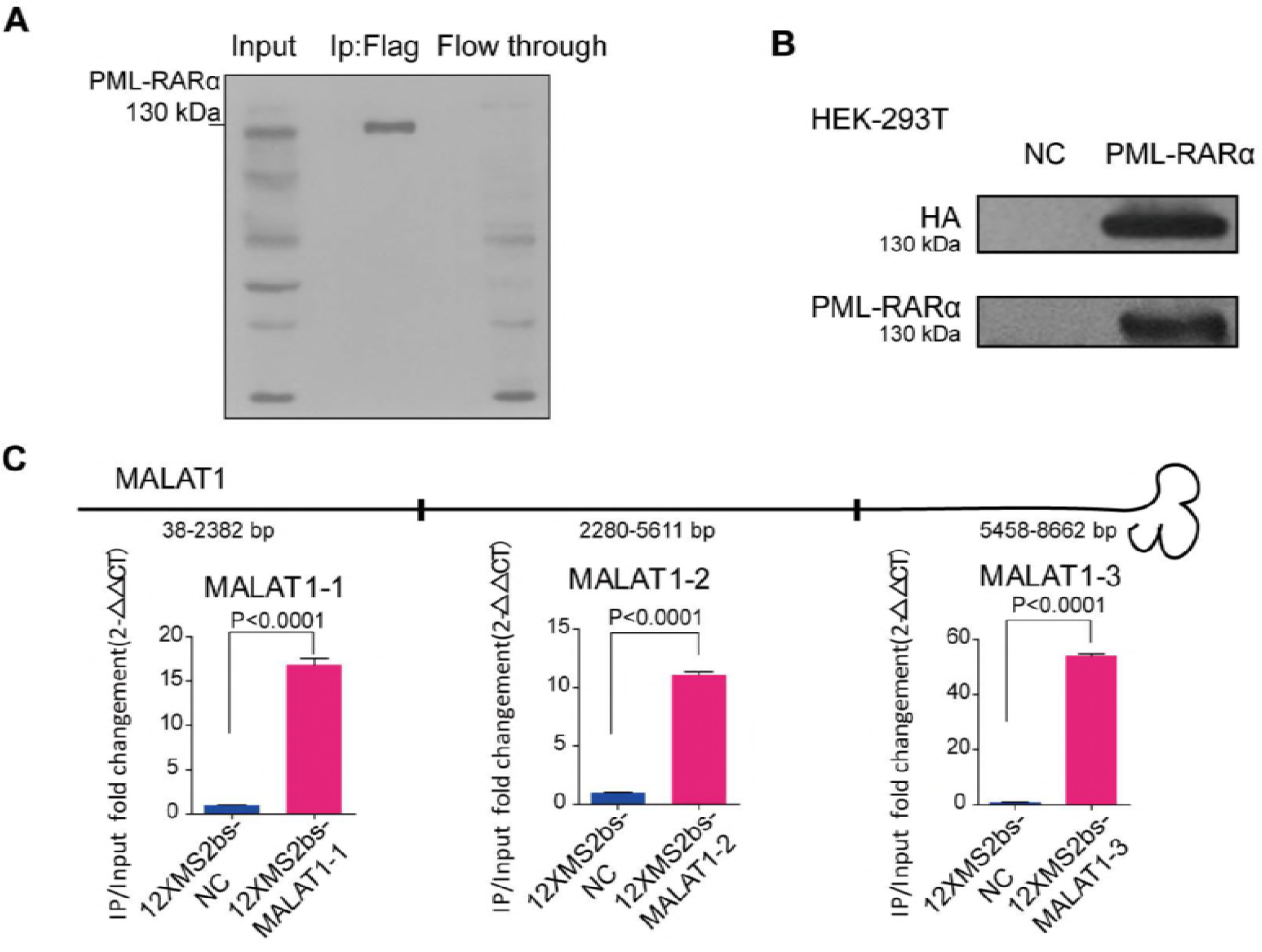
Over-expressed PML-RARA regulates cellular differentiation. **A**. Immunoprecipitation experiments were carried out in NB4 cells lentivirally infected with a Flag-tagged PML-RARA PCDH construct, and the subsequent Western blot assay showed that the specific PML-RARA was observed. **B**. The same results for Western blot using antibodies for HA or RARA showed that compared with the negative control, HA-tagged PML-RARA was also overexpressed in liposome-transfected HEK-293T cells. **C**. Schematic showing MALAT1 sections in MS2-Trap assays. qRT-PCR results showed that MALAT1 was much more enriched in 12×MS2bs-MALAT1 samples than in negative controls 12×MS2bs-NC after the immunoprecipitation of Flag-tagged MS2. Data are shown as the means ± s.e.m.; n = 3 independent experiments.

**Figure EV2.**
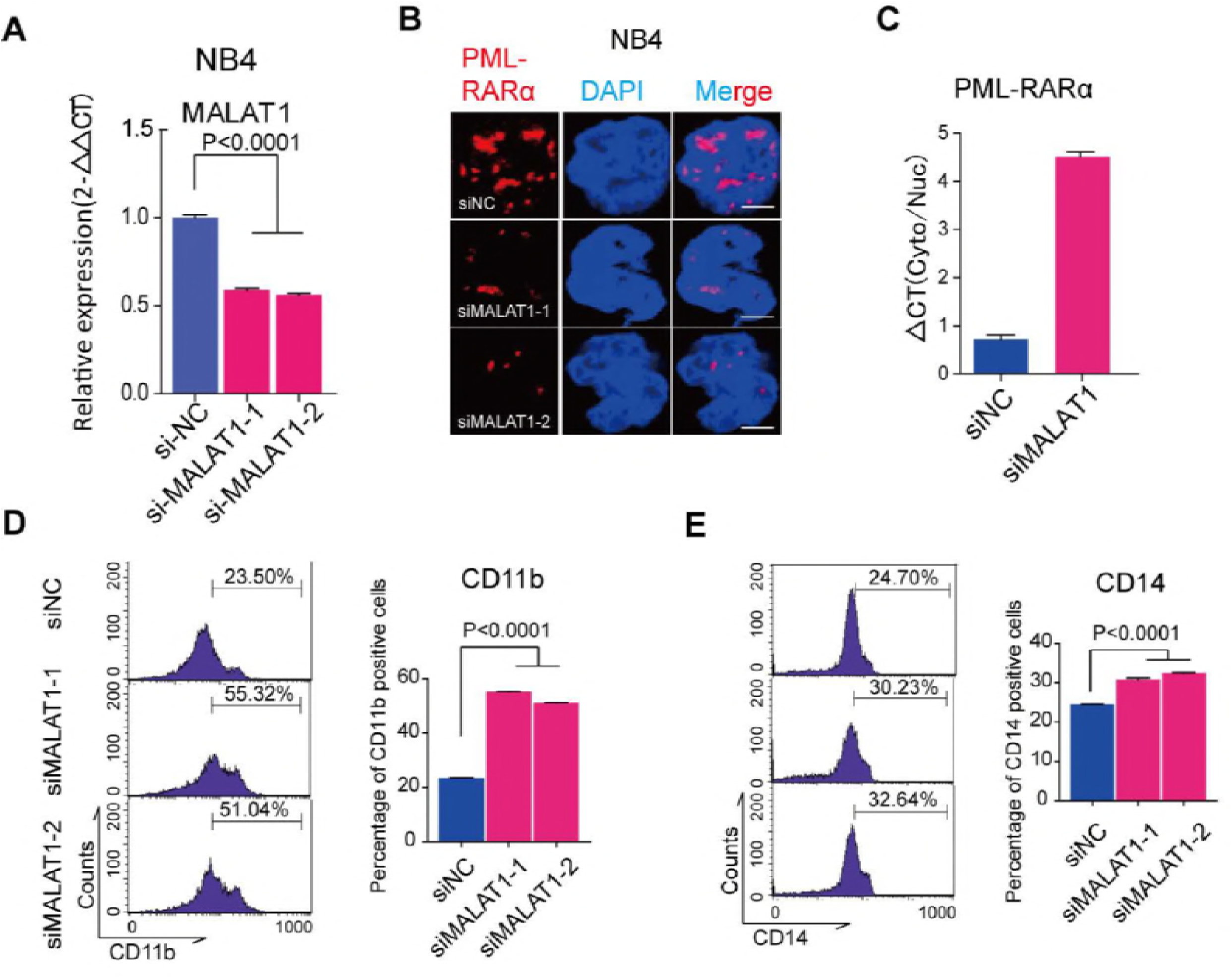
Reduction of MALAT1 promoted cellular differentiation by inhibiting the mRNA export process of fusion proteins. **A**. qRT-PCR measurement of MALAT1 levels after suppressing MALAT1 by two siRNAs with different primers of MALAT1 in NB4 cells. Data are shown as the mean ± s.e.m.; n = 3 independent experiments. **B**. Immunofluorescence analysis of PML-RARA in NB4 cells and representative images showing that the dots indicating PML-RARA were markedly decreased after knockdown of MALAT1 compared with those in controls. Scale bar represents 4 μm. **C**. qRT-PCR delta CT analysis of PML-RARA mRNA levels in the cytoplasm versus the Nucleus. Data are shown as the means ± s.e.m.; n = 3 independent experiments. **D,E**. Cellular differentiation markers CD11b and CD14 were detected by Flow cytometry. The results showed that compared to the negative controls, both markers were increased significantly when MALAT1 was down-regulated. The data were analyzed from at least three independent experiments and are shown as the means ± s.e.m.

**Figure EV3.**
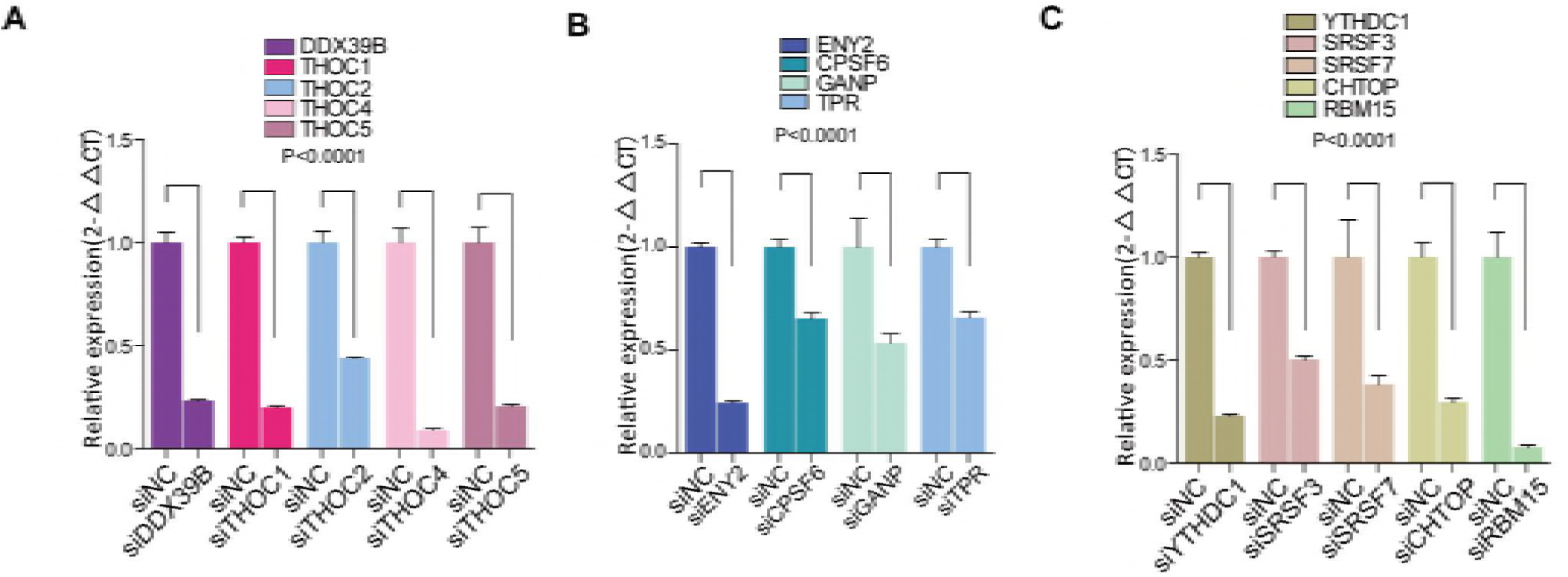
Screening of mRNA export regulatory proteins required for fusion gene transport from the nucleus to the cytoplasm. **A-C**. Measuring the effects of knocking down mRNA export-related proteins by siRNAs using qRT-PCR results normalized to the reference gene GAPDH. Data are shown as the means ± s.e.m.; n = 3 independent experiments.

**Figure EV4.**
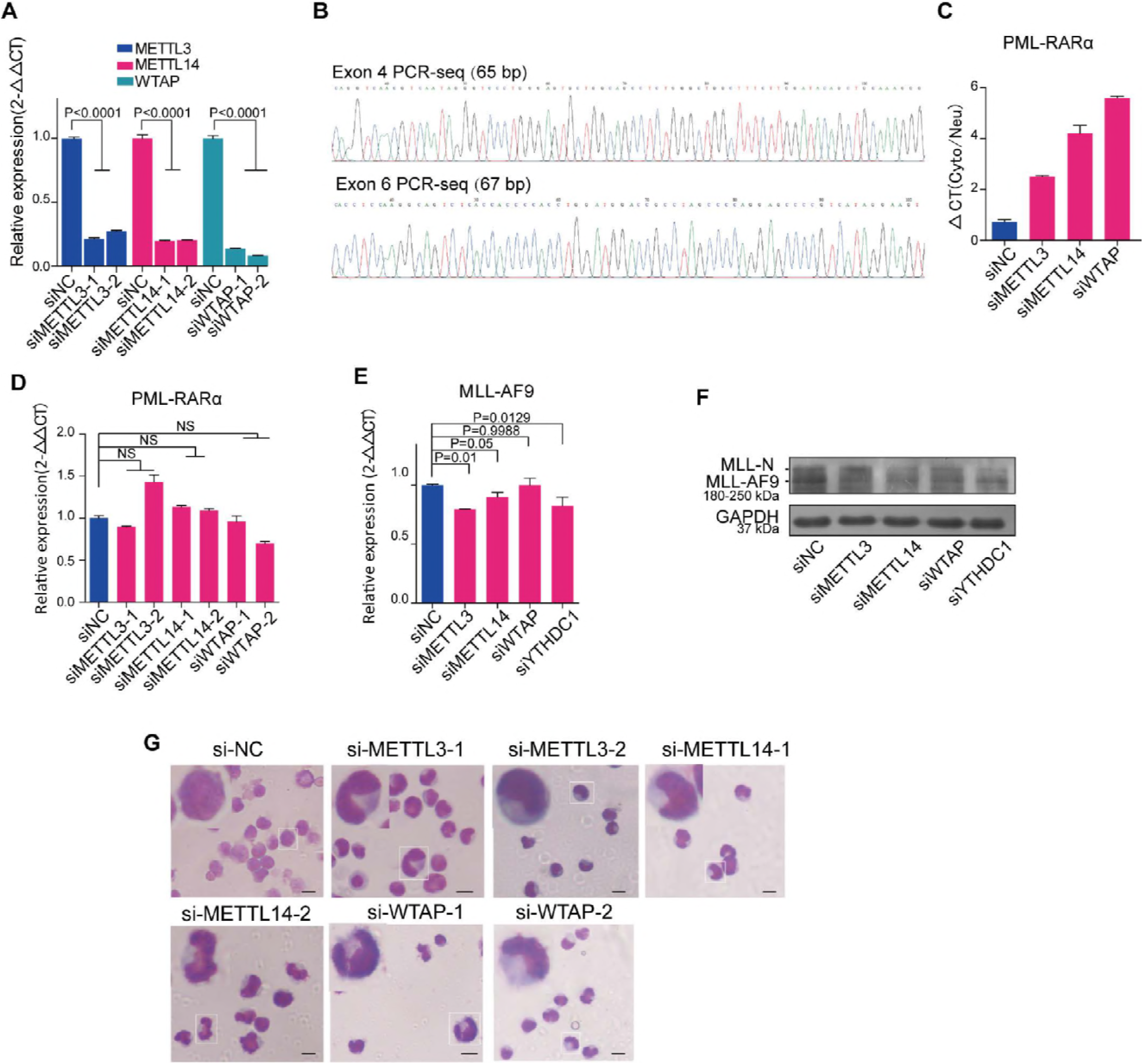
MALAT1-regulated m6A levels are responsible for fusion gene export and cellular differentiation. **A**. Measuring the effect of knocking down m6A methyltransferases by siRNAs using qRT-PCR normalized to the reference gene GAPDH. Data are shown as the means ± s.e.m.; n = 3 independent experiments. **B**. Sequencing results of MeRIP-PCR for PML exon 4 and exon 6. **C**. The qRT-PCR delta CT analysis of PML-RARA mRNA levels in the cytoplasm versus the nucleus in NB4 cells after decreasing m6A methyltransferase levels. Data are shown as the means ± s.e.m.; n = 3 independent experiments. **D,E**. PML-RARA and MLL-AF9 mRNA detection by qRT-PCR after inhibiting m6A levels or YTHDC1 in NB4 cells and moml13 cells, respectively. **F**. The protein levels of MLL-AF9 were monitored via Western blot after knocking down m6A methyltransferases and YTHDC1 by siRNAs. Three independent experiments were carried out. **G**. Wright-Giemsa staining showed that cellular differentiation was significantly promoted after reducing m6A levels in NB4 cells. The nuclear shape of these cells was altered from a round to a lobulated shape. The experiments were performed independently at least three times. Scale bar represents 10 μm.

**Figure EV5.**
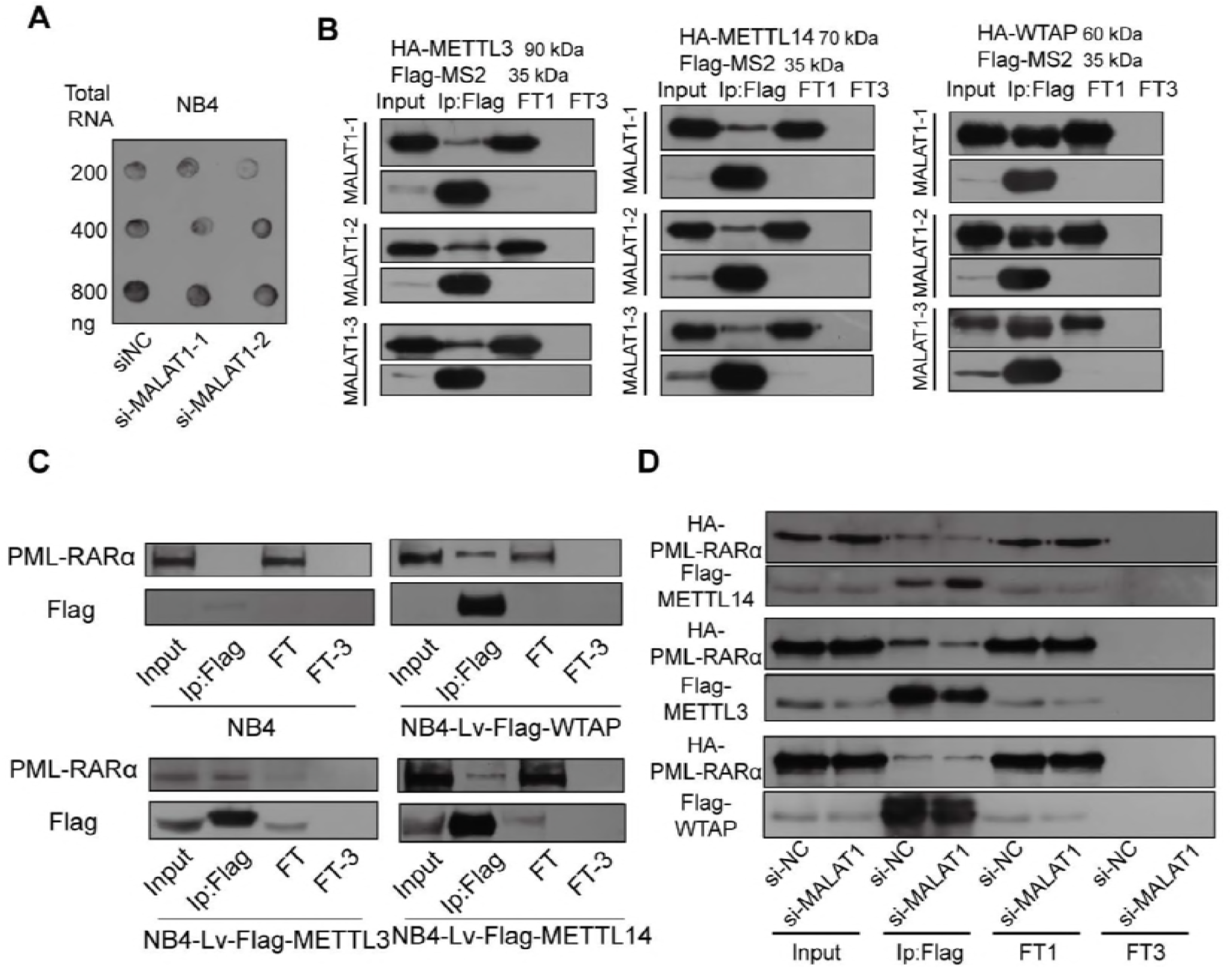
MALAT1 is involved in fusion protein and m6A methyltransferase complex formation and regulates their interactions. **A**. Dot blot assays were performed to determine cellular m6A levels after knockdown of MALAT1. **B**. MS2-Trap assays were carried out to test the interactions of METTL3, 14 and WTAP with the three sections of MALAT1. Flow through (FT1) and flow through after washing three times (FT3) were loaded as controls. **C**. Western blot analysis for PML-RARA following the immunoprecipitation of specific m6A methyltransferases in the lentiviral-infected NB4 cells.. Flow through (FT1) and flow through after washing three times (FT3) were loaded as controls. **D**. Co-immunoprecipitation of HA-tagged PML-RARA with Flag-tagged METTL3, 14 and WTAP in HEK-293T cells with MALAT1 knockdown by siRNA. Flow through (FT1) and flow through after washing three times (FT3) samples were loaded as controls.

**Figure EV6.**
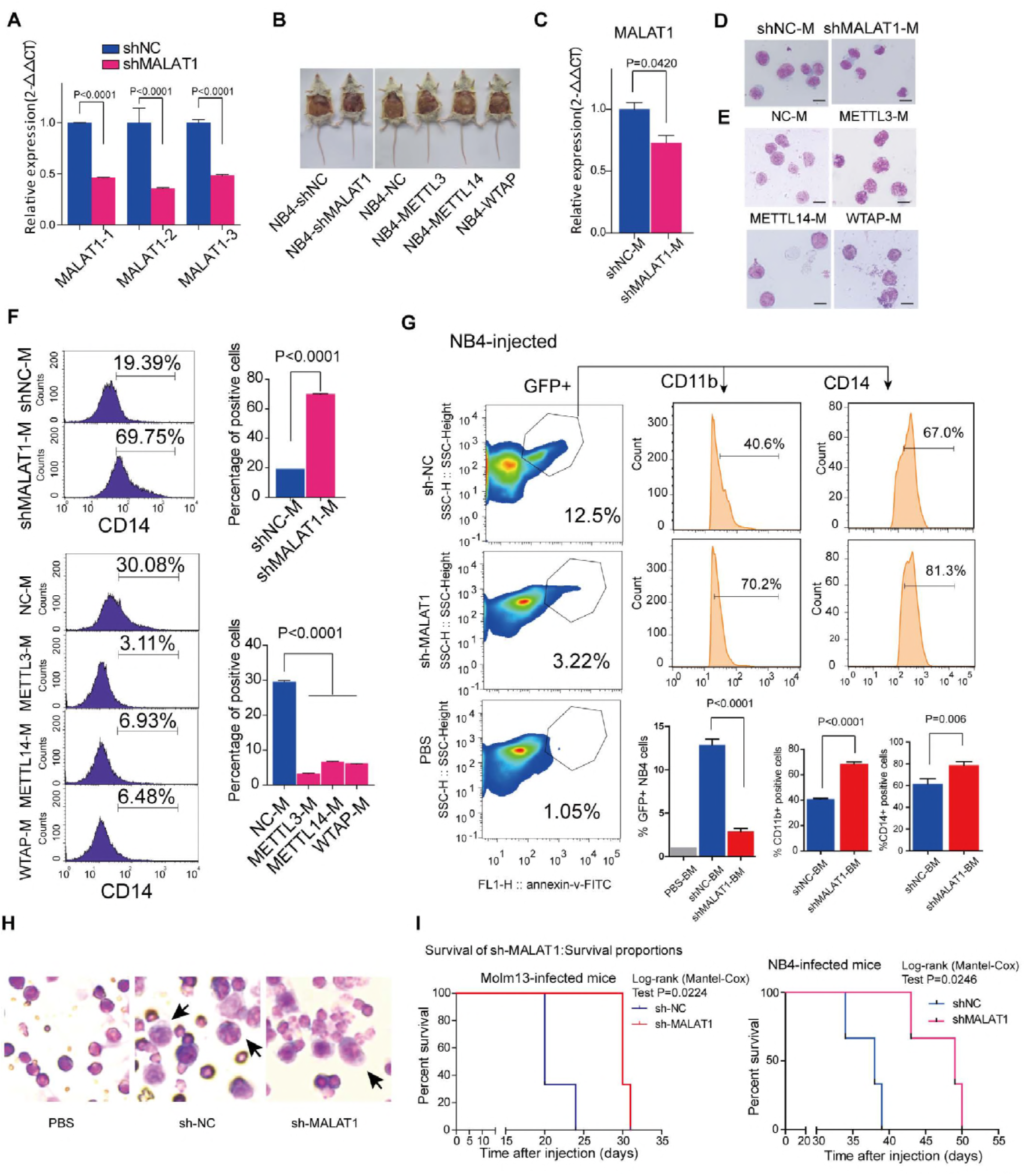
MALAT1 functions in cellular differentiation via m6A during malignant hematopoiesis in vivo. **A**. Determining the effect of knocking down MALAT1 in NB4 cells infected by lentiviral constructs carrying shRNA-targeted MALAT1 using qRT-PCR normalized to GAPDH. Data are shown as the means ± s.e.m.; n = 3 independent experiments. **B**. Representative images of NOD-SCID mice with ascites fluid injected by human NB4 cells that were suppressed for MALAT1 or enhanced m6A methyltransferases. **C**. Testing the effects of knocking down MALAT1 of ascites fluid of NOD-SCID mice infected by lentiviral constructs carrying shRNA targeting MALAT1 using the methods of qRT-PCR normalized to reference gene GAPDH. Data are shown as the means ± s.e.m.; n = 3 independent experiments. **D,E**. Wright-Giemsa assays were carried out to verify the karyotypes of murine ascites fluid cells after knocking down MALAT1 or overexpressing METTL3, 14 or WTAP in human NB4 cells. Scale bar represents 10 μm. **F**. Differentiation marker CD14 of NOD-SCID murine ascites fluid was analyzed by flow cytometry after altering MALAT1 expression or m6A levels. Data are shown as the mean ± s.e.m.; n = 3 independent experiments. **G**. NOD-SCID mice intravenously injected by NB4 cells were analyzed by flow cytometery after reducing MALAT1. 3 mice each group, the mice injected the PBS as the negative control (n=1).Analyzing the ratio of CD11b and CD14 positive cells in BM of mice injected by NB4 cells. Data are shown as the mean ± s.e.m.; n = 3 independent experiments. **H**. Wright-Giemsa assays were carried out to verify the karyotype changed to lobulated in human shMALAT1 molm13 cells transplanted into BM. Scale bar represents 10 μm. **I**. Survival curve of mice after intravenous injection of Molm13 and NB4 cells. Data were shown that the survival time of mice injected NB4 and Molm13 shMALAT1 cells significantly was longer than that in the control NB4 and Molm13 shNC cells. 3 mice each group.

**Table EV1.**
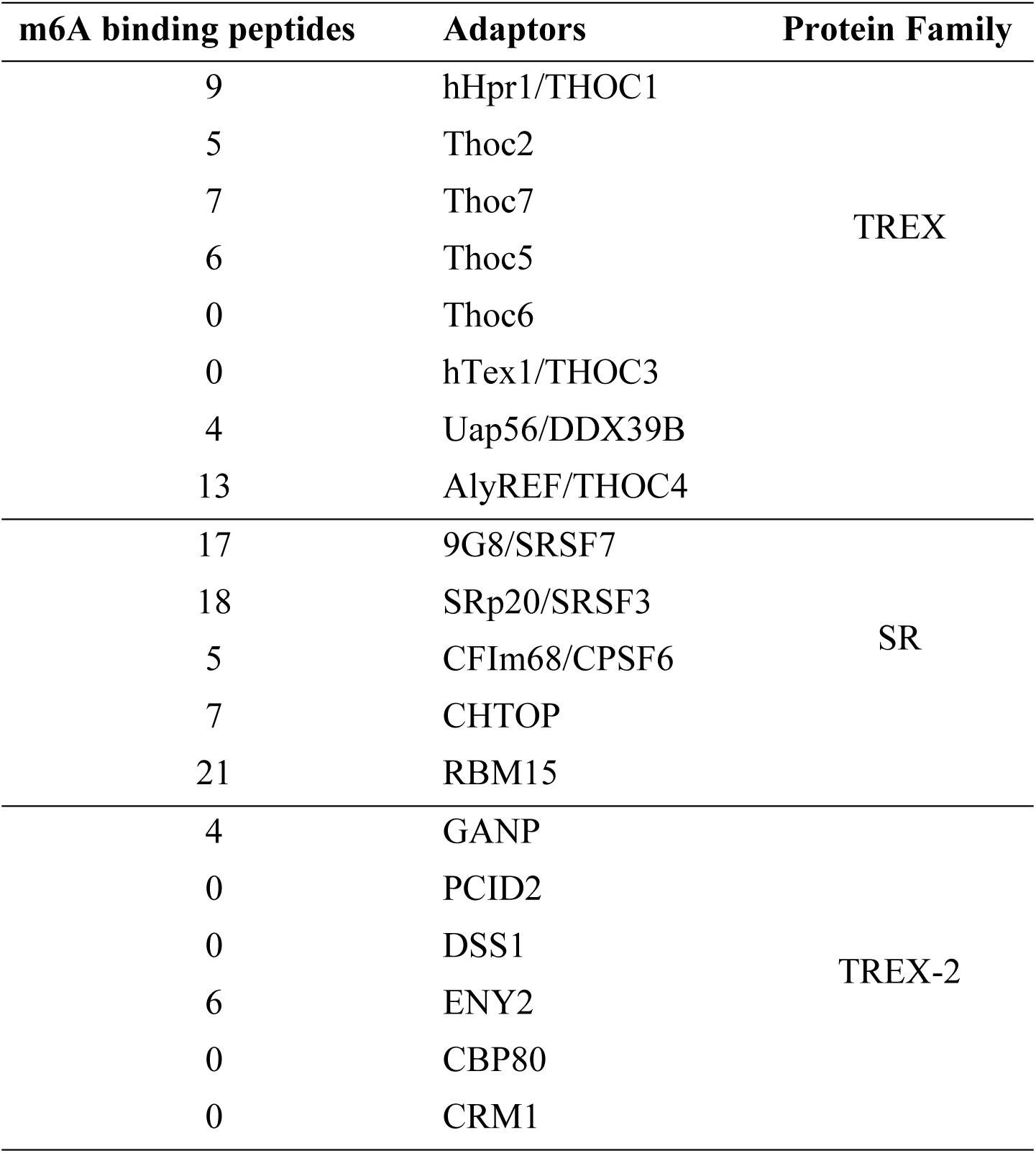
Adaptor proteins regulating mRNA export.

**Table EV2.** The predicted m6A motif located at PML-RARα mRNA.

| M6A<br>methylation | conserved<br>binding<br>sites | motif of<br>PML<br>fusion<br>exon 4<br>up and<br>down<br>1kb | motif of<br>PML<br>fusion<br>Exon 4 | motif of<br>PML<br>fusion<br>exon 5<br>up<br>1kb | motif of<br>PML<br>fusion<br>Exon 5 | motif of<br>PML<br>fusion<br>exon 6<br>up and<br>down<br>1kb | motif of<br>PML<br>fusion<br>Exon 6 | motif of<br>RARA<br>fusion<br>exon 3<br>up and<br>down<br>1kb | motif of<br>RARA<br>fusion<br>exon 3 |
| --- | --- | --- | --- | --- | --- | --- | --- | --- | --- |
| METTL14 | GGACT | 4 | 0 | 2 | 0 | 1 | 0 | 2 | 0 |
|  | GGACC | 2 | 1 | 1 | 0 | 1 | 1 | 0 | 0 |
|  | GGACA | 4 | 0 | 1 | 0 | 4 | 0 | 4 | 1 |
| METTL3 | GAGGAC | 0 | 0 | 1 | 0 | 0 | 0 | 0 | 0 |
|  | GTGGAC | 0 | 0 | 0 | 0 | 2 | 0 | 0 | 0 |
|  | GCGGAC | 0 | 0 | 0 | 0 | 0 | 0 | 0 | 0 |
|  | CAGGAC | 1 | 0 | 0 | 0 | 0 | 0 | 1 | 1 |
|  | CTGGAC | 5 | 0 | 1 | 0 | 1 | 0 | 0 | 0 |
| WTAP | GACTCT | 1 | 0 | 0 | 0 | 0 | 0 | 2 | 0 |
|  | GGCTCT | 1 | 0 | 0 | 0 | 2 | 0 | 2 | 0 |
|  | GACTGT | 1 | 0 | 0 | 0 | 0 | 0 | 0 | 0 |
| **Total** |  | **19** | **1** | **6** | **0** | **11** | **1** | **11** | **2** |

**Table EV3.**
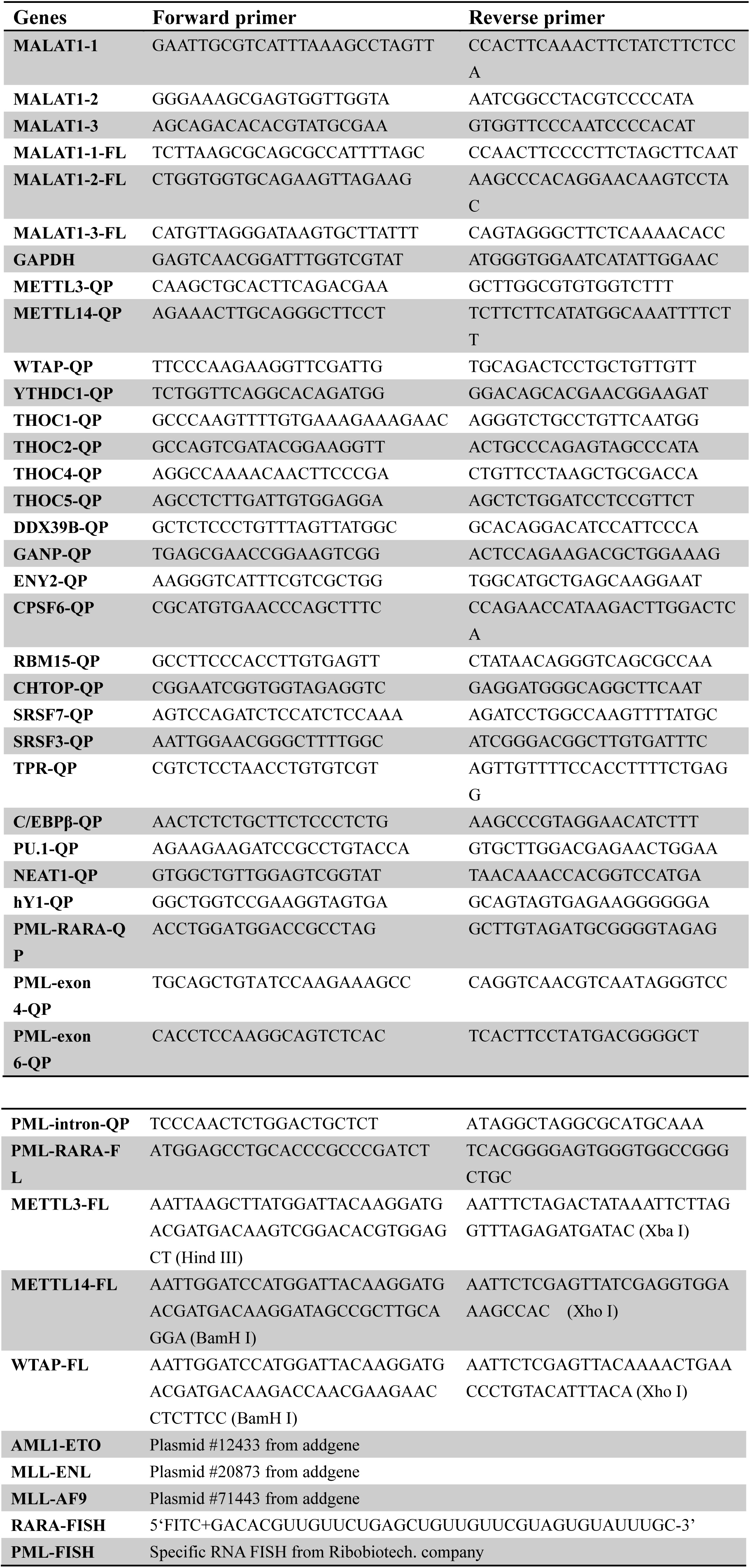
Primers used in qRT-PCR (QP) and full-length (FL) cDNA amplification.

**Table EV4.**
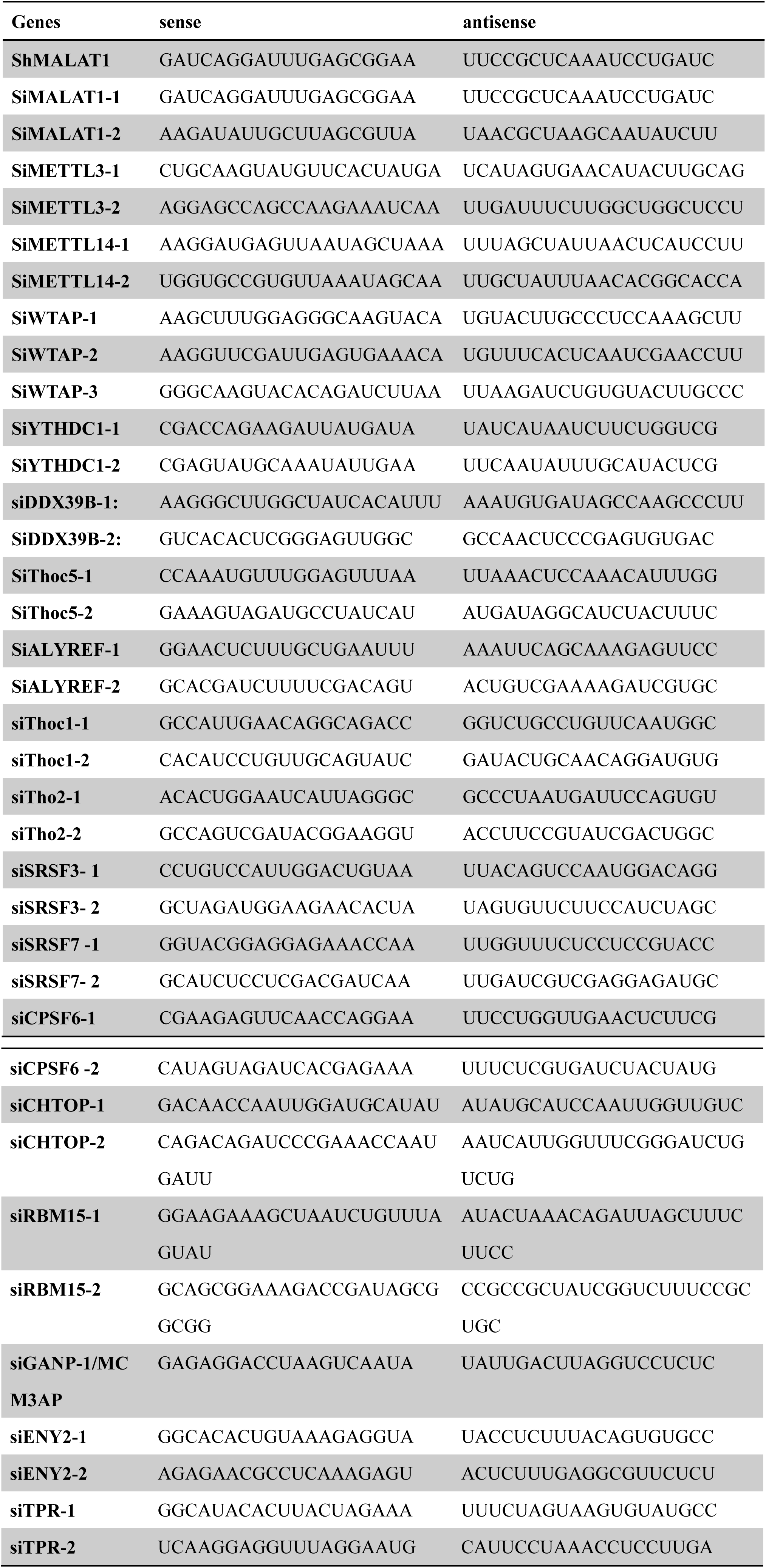
siRNA or shRNA sequences for target genes.

**Table EV5.** Antibodies used in WB and IF.

| **Antibody** | **Company** | **Catalog** |
| --- | --- | --- |
| **anti-METTL3 antibody** | Proteintech | 15073-1-AP |
| **anti-METTL14 antibody** | Proteintech | 26158-1-AP |
| **anti-WTAP antibody** | Proteintech | 60188-1-Ig |
| **anti-Flag (mouse) HRP-conjugated antibody** | Sigma | A8592 |
| **polyclonal rabbit anti GAPDH) antibody** | Sigma-Aldrich | G9545 |
| **anti-Flag (rabbit) antibody** | Proteintech | 20543-1-AP |
| **anti-RARA antibody** | Santa cruz | sc-550 |
| **anti-PML antibody** | Santa cruz | sc-966 |
| **anti-HA (mouse) antibody** | Sigma | H3663 |
| **anti-HA (rabbit) antibody** | CST | C29F4 |
| **anti-MLL antibody** | Bethyl | A300-086A-3 |
| **anti-SC35 antibody** | Abcam | Ab11826 |
| **anti-SF2 antibody** | Santacruz | Sc-33652 |
| **anti-mouse HRP-conjugated secondary antibody** | Sigma Aldrich | A3682 |
| **anti-rabbit HRP-conjugated secondary antibody** | Sigma Aldrich | A9542 |
| **anti-CD11b-APC** | BD | 550019 |
| **anti-CD14-APC** | BD | 555399 |
| **anti-m6A** | Abcam | Ab151230 |
| **AlexaFluor 594-conjugated secondary antibodies(mouse)** | Invitrogen | A11005 |
| **AlexaFluor 594-conjugated secondary antibodies(rabbit)** | Invitrogen | A21207 |
| **AlexaFluor 488-conjugated secondary antibodies(mouse)** | Invitrogen | A11001 |

